# Enhancing Mask Activity in Dopaminergic Neurons Extends Lifespan in Flies

**DOI:** 10.1101/2020.06.15.153056

**Authors:** Xiaolin Tian

## Abstract

Dopaminergic neurons (DANs) are essential modulators for brain functions involving memory formation, reward processing, and decision making. Here I demonstrate a novel and important function of the DANs in regulating aging and longevity. Overexpressing the putative scaffolding protein Mask in two small groups of DANs in flies can significantly extend the lifespan in flies and sustain adult locomotor and fecundity at old ages. Such a Mask-induced beneficial effect requires dopaminergic transmission but cannot be recapitulated by elevating dopamine production in the DANs. Independent activation of Gα_s_ in the same two groups of DANs via the drug-inducible DREADD system also extends fly lifespan, further indicating the connection of specific DANs to aging control. The Mask-induced lifespan extension appears to depend on the function of Mask to regulate microtubule (MT) stability. A structure-function analysis demonstrated that the ankyrin repeats domain in the Mask protein is both necessary for regulating MT stability (when expressed in muscles and motor neurons) and sufficient to prolong longevity (when expressed in the two groups of DANs). Furthermore, DAN-specific overexpression of Unc-104 or knockdown of p150^Glued^, two independent interventions previously shown to impact MTs dynamics, also extend lifespan in flies. Together these data demonstrated a novel DANs-dependent mechanism that, upon neuronal cell-type specific tuning of MT dynamics, modulates systemic aging and longevity in flies.

## Introduction

The complex process of aging has a reciprocal interaction with the functions of the brain. This connection was indicated by studies of the neuronal specific genes and their ability to affect lifespan in animals^1^, the sensory functions of the nervous system to sense and coordinate the responses to environmental cues that lead to impact on aging and longevity in animals^2–10^, and the neuroendocrine system that modulates aging and longevity^11–15^. An recently emerging theme for the brain-driven regulation on aging highlights the change of longevity induced by neuronal cell-type-specific modulations. Recent studies showed Forkhead family transcription factors that are differentially expressed in the glia and neurons each enable distinct capacities to affect longevity^16^. Specific neuronal types such as the serotonergic neurons in both worms^17,18^ and flies^19–21^ regulate aging through targeting different physiological functions in the peripheral tissues.

Dopaminergic neurons (DANs), a crucial modulatory system in the brain^22,23^, have also been implicated in aging regulation. Limited studies on this topic yielded controversial results: on one hand, the function of the dopaminergic system in regulating reward-feeding, food intake and energy balance^24–26^ imply an intuitive link to the control of aging. The results of a few previous studies support such a link: in rodents, increasing dopamine level leads to extended longevity^27,28^. In flies, polymorphism in *dopa decarboxylase* (*Ddc*), an enzyme required for dopamine biogenesis, correlates with variations in longevities in fly populations^29^. In worms, circumventing the ER unfolded protein stress in the DANs can moderately extend lifespan^18^. On the other hand, loss of DANs or dopamine production in the brain does not seem to affect longevity. Depletion of dopamine production in the fly brains^30^ or in worms^31^ does not affect lifespans in either organisms. The life expectancy of human patients with Parkinson’s disease (PD) who lose their DA neurons in the substantia nigra is similar to the normal population^32^. Studies in human and rodent also showed that chronically and globally increasing dopamine levels led to adverse effects^33^. For example, long-term supplementation of L-DOPA together with deprenyl at low dosages cause an increased death rate in patients at an early stage of Parkinson’s disease^34^. Therefore, it remains elusive as whether and how the dopamine system can actively affect aging.

My studies on a putative scaffolding protein, Mask, provided new evidence supporting a function of DANs in regulating aging and longevity. Mask was previously implicated in diverse signaling pathways and cellular processes. In mitotic cells, Mask acts as a modulator of the receptor tyrosine kinase signaling in flies^35^, a co-factor of the Hippo pathway effector Yorkie^36,37^ and a positive regulator of the JAK/STAT pathway^38^. Our studies of Mask in post-mitotic cells, such as muscles and neurons, demonstrated that it modulates mitochondrial morphology^39^, autophagy^40^, microtubule (MT) dynamics and synaptic morphology^41^. In this study, I show that overexpressing Mask in two small groups of DANs in the fly brain significantly extends the lifespan of these flies, and this DAN-mediated effect can be independently recapitulated by moderately activating Gα_s_ in the two groups of DANs. I also provide multiple pieces of evidence to support a model that enhanced MT dynamics in the DANs is a primary contributing factors conveying dopaminergic-dependent lifespan extension. All these results together point to the notion that MT stability and dynamics in specific DANs is a targetable process that can be tuned to impact aging.

## Results

### Small subsets of the DANs regulates longevity in flies

Our previous studies on Mask uncovered diverse functions of this putative scaffolding protein in the post-mitotic cells including a strong protective effect against neurodegeneration induced by toxic aggregation-prone proteins^42^. In the continued investigation seeking to understand Mask’s functions in neurons, I uncovered an interesting gain-of-function effect of Mask in the DANs. Overexpressing Mask in DANs (driven by pan-dopaminergic TH-Gal4 driver) leads to a ~40% increase in lifespan in flies (median lifespan in male flies increased from ~76 days to ~104 days) (Fig. S1). Overexpressing Mask in either the entire body or the nervous system showed no effects on the lifespan (Fig. S2), suggesting that other neuronal types potentially counteract the effects of DNAs on aging. Dopaminergic neurons in the adult fly brain cluster into eight groups^43^, and using three well-characterized Gal4 drivers that together show largely non-overlapping expression in most of the dopaminergic neurons in the fly brains^44–46^, I tested whether all or only subsets of dopaminergic neurons are responsible for the Mask-mediated lifespan-extension function. Overexpressing Mask in these three groups of dopaminergic neurons yielded distinct results on longevity. In the PAM dopaminergic neurons marked by the 0273-Gal4 driver, overexpressing Mask does not alter the lifespan (Fig. 1A); however, expressing Mask driven by either the TH-C’- or TH-D’-Gal4 drivers significantly extends the lifespan – the median lifespan in male flies increased from 64 days to 96 days, and from 63 days to 86 days, respectively (Fig. 1 B&C).

**Figure 1.**
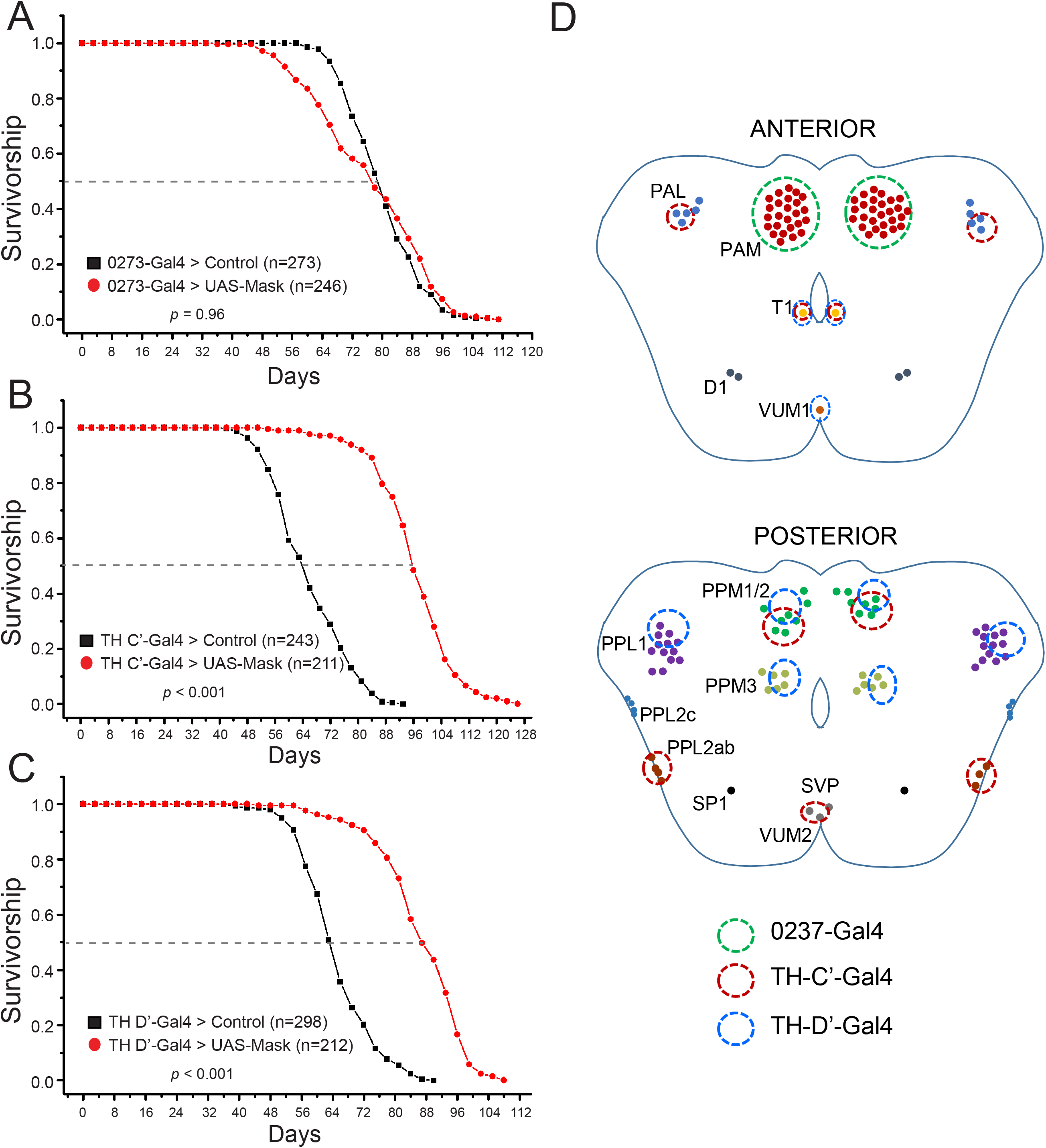
Overexpressing Mask in the TH-C’ or TH-D’ DANs Extends Lifespan. Survivorship curves of control flies and flies overexpressing Mask under the control of A) 0273-Gal4, B) TH-C’-Gal4, and C) TH-D’-Gal4 drivers. D) Schematic of expression patterns of 0273-, TH-C’– and TH-D’-Gal4 drivers.

Although the dopaminergic system provides essential modulation on various behaviors and physiological functions, flies devoid of dopamine in their brains^30^ and worms lacking the rate-limiting enzymes for dopamine synthesis both live a normal lifespan^31^. Consistently, blocking neuronal activity in the TH-C’ or TH-D’ dopaminergic neurons by the DREADD-Di^47^ does not affect the lifespan in flies (Fig. S3). Together, these results suggest that dopamine systems may be dispensable for the mechanisms that drive the normal aging process. Overexpressing Mask in the specific dopaminergic neurons possibly induces a gain-of-function cellular effect, which consequently confers a beneficial outcome on aging and longevity. Before investigating the underlying mechanisms induced by Mask, I first set out to determine whether activation of the DANs through approaches independent to Mask overexpression could also yield similar effects on longevity. I found that continuously full activation of the TH-C’ or TH-D’ dopaminergic neurons via the thermosensitive cation channel TrpA1 shortened the lifespan in the male flies and had no effect on the female flies (Fig. S4). However, moderate activation of these DANs induced lifespan extension in the flies (Fig. 2). The inducible DREADD-Ds receptor when expressed in neurons permits dose-dependent neuronal modulation^47^. Feeding 1 μM Clozapine N-oxide (CNO) (a concentration significantly below the full activation dose^47^) to flies expressing this engineered Gα_s_ receptor in their TH-C’ or TH-D’ neurons induces prominent and consistent lifespan extension (Fig. 2). These results suggest that moderate modulation of these DANs independent of Mask expression is sufficient to enhance longevity in flies, further confirm the potentials of the DANs to impact lifespan.

**Figure 2.**
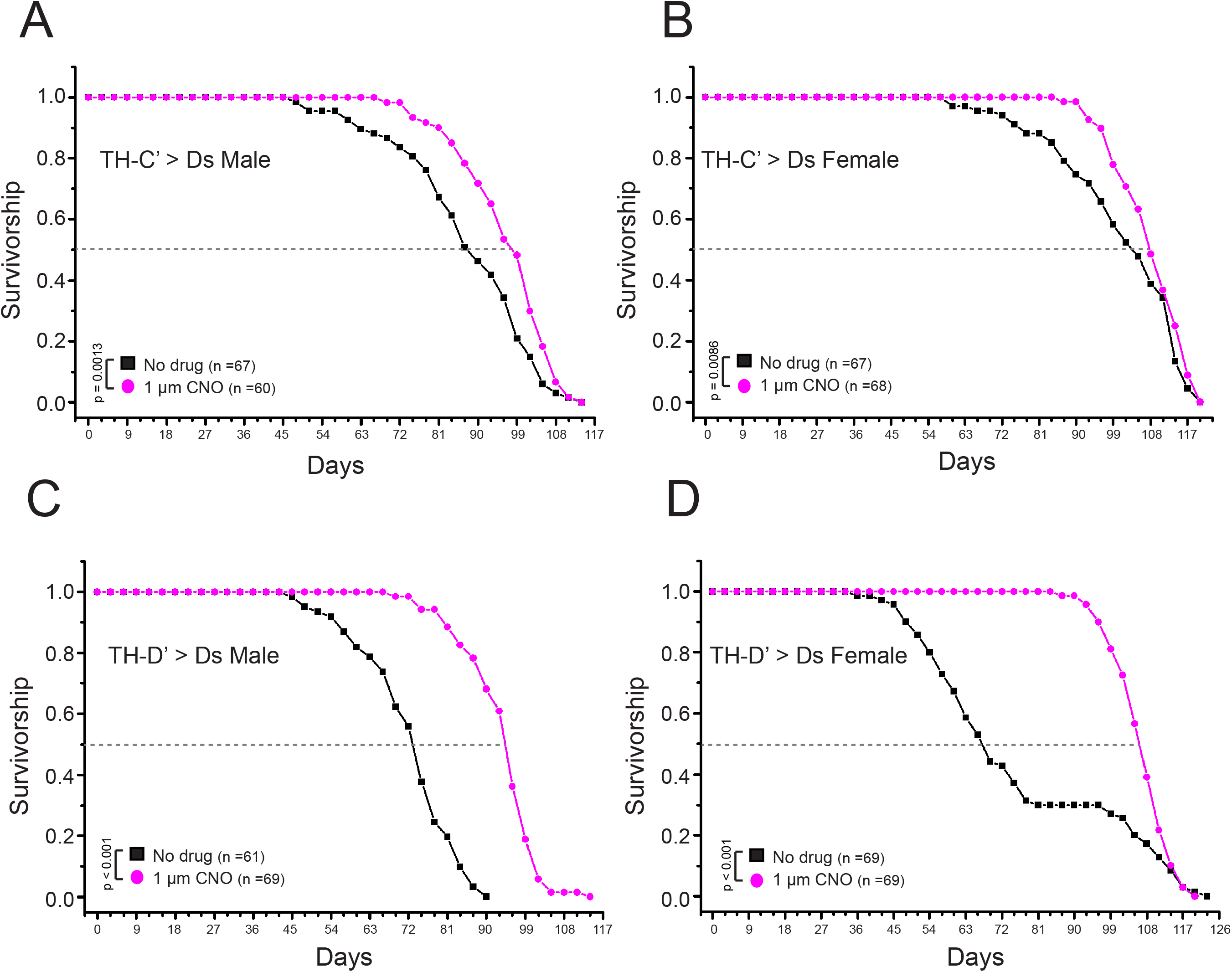
Moderately activating the ectopically expressed DREADD system in the TH-C’ or TH-D’ DANs Extends Lifespan in flies. Survivorship curves of male and female adult flies expressing the DREADD Gα_s_ receptor and raised on food containing DMSO only (No Drug) or 1uM Clozapine N-oxide (CNO). A, B) moderately activating the DREADD-Ds receptor in the TH-C’ neurons leads to extension of lifespan in flies. The median lifespan increased from ~87 days to ~98 days (~12.6% increase) in the male flies and from ~95 days to ~108 days (~12.6% increase) in the female flies. C, D) moderate activation of the DREADD-Ds receptor expressed in the TH-D’ neurons leads to significant extension of lifespan in flies. The median lifespan increased from ~73 days to ~95 days (~30% increase) in the male flies and from ~67 days to ~107 days (~50% increase) in the female flies.

### Dopamine transmission is required to elicit the Mask-induced lifespan extension

The Tyrosine Hydroxylase (TH)-positive neurons often use dopamine as the neural transmitter to communicate with other neurons but they may also co-transmit additional transmitter^48–50^. Next, I asked whether dopamine transmission is required in mediating the lifespan extension induced by Mask-overexpression in the DANs. If dopamine transmission is needed, reducing dopamine signaling in these neurons would suppress the strength of the neuronal communication which in turn may hinder the ability of Mask-overexpression to extend lifespan. Four genes are known to be essential for dopamine transmission: two enzymes responsible for dopamine synthesis - Tyrosine Hydroxylase (TH) and Dopa Decarboxylase (Ddc); and two transporters for dopamine - dopamine transporter (DAT) that controls dopamine reuptake, and vesicular monoamine transporter (VMAT) that mediates synaptic vesicular loading of dopamine. Genetically heterozygosity that reduces gene dosages of either of these four genes was introduced to the Mask-overexpression paradigm to test whether dopamine transmission is necessary for the lifespan extension. The results show that reducing the dosage of each of these four genes can blunt the degree of lifespan extension in flies that overexpress Mask in the TH-C’ or TH-D’ neurons compared to the wild type background (Fig. 3). These results support the notion that the dopamine system is necessary for Mask to induce lifespan extension.

**Figure 3.**
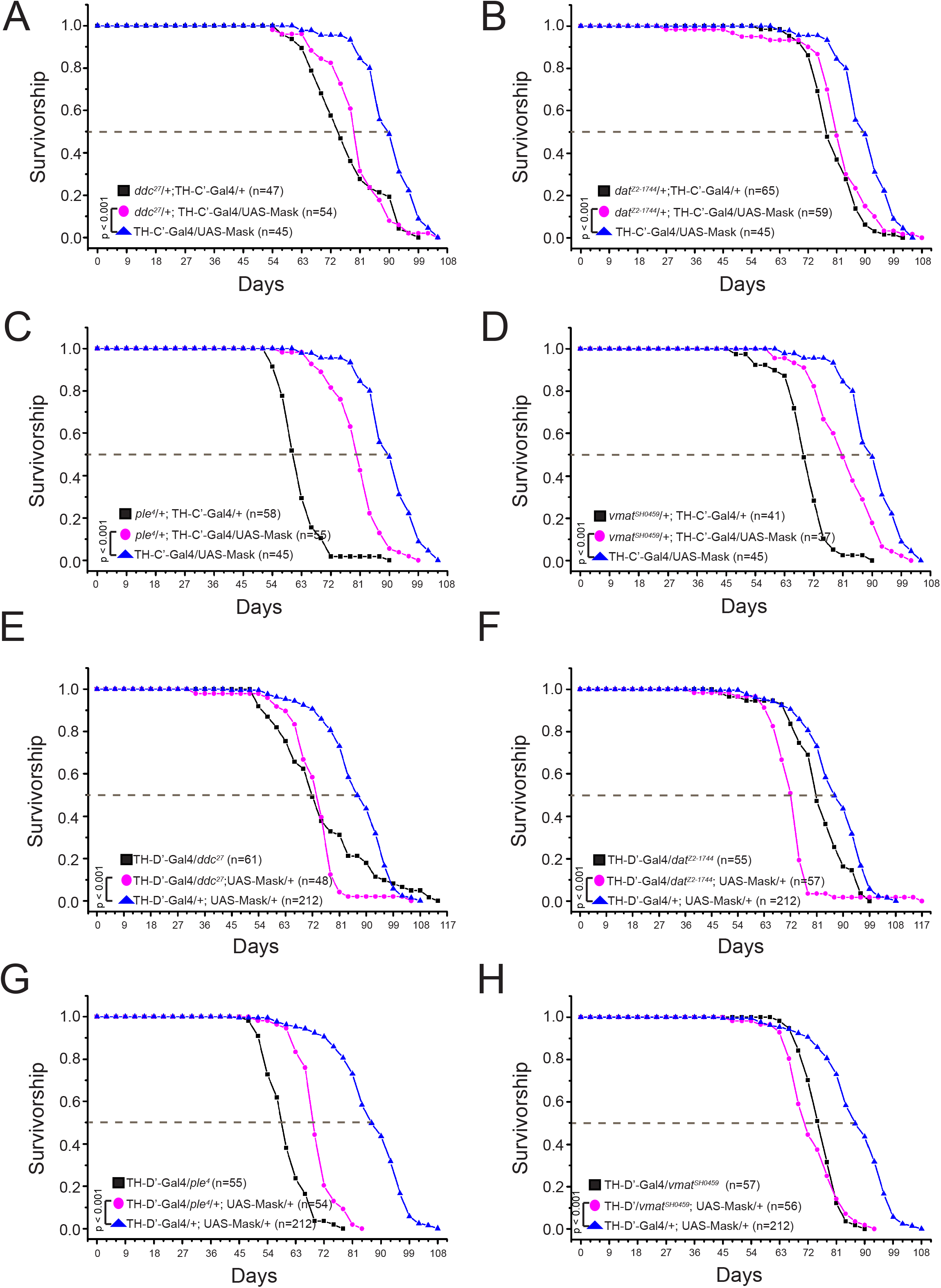
Reducing Dopamine Pathway Activity Blunts the Lifespan Extension effects induced by Overexpressing Mask in the TH-C’ or TH-D’ DANs. Decreasing dopaminergic pathway activity was achieved by genetically reducing the dosage of the four genes that are essential for dopamine transmission: two enzymes responsible for dopamine synthesis - TyrosineHydroxylase (TH) and Dopa Decarboxylase (Ddc); and two transporters for dopamine - dopamine transporter (DAT) that controls dopamine reuptake, and vesicular monoamine transporter (VMAT) that mediates synaptic vesicular loading of dopamine. Four loss of function alleles, *ple^4^* (TH), *ddc^27^*, *dat^Z2-1744^* or *vmat^SH0459^,* were introduced to Mask-overexpressing paradigms to generate a heterozygous background of each gene. The survivorship curves of male flies of the following genotypes are shown: A) Overexpressing Mask in the TH-C’ DA neurons in wild type or in the *ddc^27^/+* heterozygous backgrounds; B) Overexpressing Mask in the TH-C’ DA neurons in wild type or in the *dat^Z2-1744^/*+ heterozygous backgrounds; C) Overexpressing Mask in the TH-C’ DA neurons in wild type or in the *ple^4^/*+ heterozygous backgrounds; D) Overexpressing Mask in the TH-C’ DA neurons in wild type or in the *vmat^SH04594^/*+ heterozygous backgrounds; E) Overexpressing Mask in the TH-D’ DA neurons in wild type or in the *ddc^27^/+* heterozygous background; F) Overexpressing Mask in the TH-D’ DA neurons in wild type or in the *dat^Z2-1744^/*+ heterozygous background; G) Overexpressing Mask in the TH-D’ DA neurons in wild type or in the *ple^4^/*+ heterozygous background; H) Overexpressing Mask in the TH-D’ DA neurons in wild type or in the *vmat^SH04594^/*+ heterozygous background. With all conditions, reducing gene dosage of TH, Ddc, DAT or VMAT is able to blunt the lifespan extension induced by overexpressing Mask in DANs.

I next tested whether upregulation of dopamine signaling in the TH-C’ or TH-D’ DANs is sufficient to induce lifespan extension in flies. In order to achieve the upregulation, the flies were fed with dopamine precursor L-DOPA^51^, dopamine D1 receptor agonist SKF-82958^52^, dopamine D2 receptor agonist Quinpirole^53^, or the D1 and D2 agonists together. It appeared that none of the treatments was able to extend the lifespan in flies. The lifespan of female flies fed with L-DOPA was even moderately shortened (Fig. S5). This adverse effects may be attributable to the global elevation of dopaminergic signaling, as long-term treatment with L-DOPA has been shown to cause neural toxicity in mammals^33^. To limit the intervention only to specific DANs, dopamine production was selectively increased in the TH-C’ or TH-D’ DANs via overexpressing Tyrosine Hydroxylase (TH), or Dopa Decarboxylase (Ddc) in these two groups of neurons. Again, overexpressing these two enzymes in the TH-C’ or TH-D’ DANs did not prolong the longevities in flies (Fig. 4). Similar to the L-DOPA feeding, these treatments lead to a moderate reduction in the lifespan in female flies. These results together suggest that although neurons require dopamine transmission to communicate and to induce lifespan extension, increasing dopamine production or activating dopamine signaling are not sufficient to elicit similar effects on aging and longevity induced by Mask overexpression or moderate DREADD-Ds activation in these neurons.

**Figure 4.**
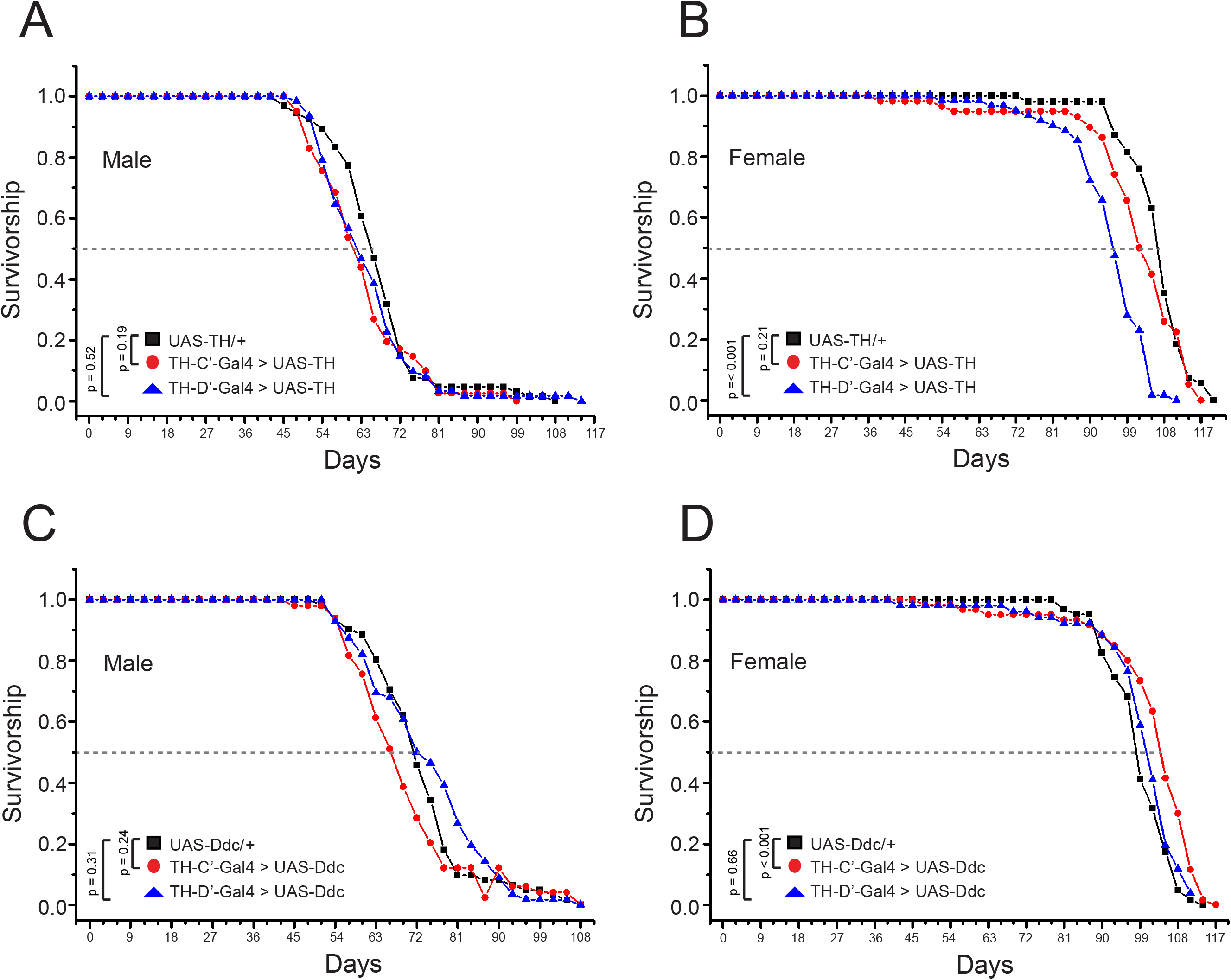
Specifically Increasing Dopamine Production in the TH-C’ or TH-D’ DANs fails to Extend Lifespan. Survivorship curves of flies expressing the rate limiting enzyme for dopamine biosynthesis tyrosine hydroxylase (TH) or Dopa Decarboxylase (Ddc) in the TH-C’ or TH-D’ neurons are shown. The longevity of A) Male and B) female flies overexpressing TH in the TH-C’ or TH-D’ DANs are recorded. This treatment does not affect the lifespan in male or female flies that overexpress TH in the TH-C’ DANs. Female flies that overexpress TH in the TH-D’ DA neurons show moderately reduced lifespan. C) Male flies overexpressing Ddc in the TH-C’ or TH-D’ DA neurons exhibit similar lifespan compared to control flies. D) Overexpressing Ddc in the TH-C’ DA neurons does not affect the lifespan. However, overexpressing Ddc in the TH-D’ DA neurons moderately extend the lifespan (median lifespan increased from 99 days in the control flies to 105 days in TH-D’-Gal4 > Ddc flies, ~6% increase).

### The long-lived Mask-overexpressing flies sustain adult locomotor and reproduction at old ages

To fully understand how aging and longevity at the systemic level are impacted by overexpressing Mask in the TH-C’ or TH-D’ neurons, I next examined age-related phenotypes including feeding, energy storage/metabolic, adult locomotor and brain insulin productions in Mask-expressing and control flies. DANs regulate feeding and food-foraging in flies^54–57^, raising the possibility that the long-lived flies that overexpress Mask in the TH-C’ or TH-D’ neurons may consume less food and therefore undertake voluntary caloric restriction, which may serve as a potential contributing factor for lifespan extension in these flies. However, the Mask-overexpressing long-lived flies consume a comparable amount of food compared to the control flies at young and middle ages (10 and 30 days old, respectively) in the Capillary Feeder assay (CAFE assay)^58^ (Fig. S7). These flies also show moderately incresed whole-body glucose and Triglyceride (TAG) levels at middle and late ages (30 and 50 days old, respectively) (Fig. S8). There is also no consistent alterations in transcription levels of the Dilp2, 3, 5, and 6 in their brains between genders and ages in the Mask-expressing flies compared to the controls (Fig. S9-12). Adult locomotor reflects overall activity and health in flies and can be quantified using the rapid iterative negative geotaxis (RING) assay^59,60^. Expressing Mask in the TH-C’ dopaminergic neurons significantly enhances the locomotor performance during the entire lifespan, and expressing Mask in the TH-D’ neurons sustain the adult locomotor at middle and late ages (Fig. 4). Altogether, these results demonstrated that the Mask-overexpressing flies live healthy and active lives throughout their entire lifespan.

Delayed reproductive maturation or reduced fecundity correlates closely with lifespan extension^61^. To test whether there is a trade-off between reproduction and lifespan in Mask-overexpressing flies, I examined the fecundity of male and female flies at different ages. No delay or reduction in reproduction in either male or female flies was detected. Instead, overexpressing Mask in the TH-C’ neurons lead to increased fertility in female flies beginning at the onset of the reproduction maturation, and the TH-D’-overexpressing flies showed a significantly higher level of fecundity at older ages (Fig. 5). Clearly, Mask-overexpressing flies show a trait of extended lifespan with no cost of reproduction; and along with a number of other previous studies of longevity-related mutant flies^62–65^, this trait contradicts the long acknowledged disposable soma theory of aging.

**Figure 5.**
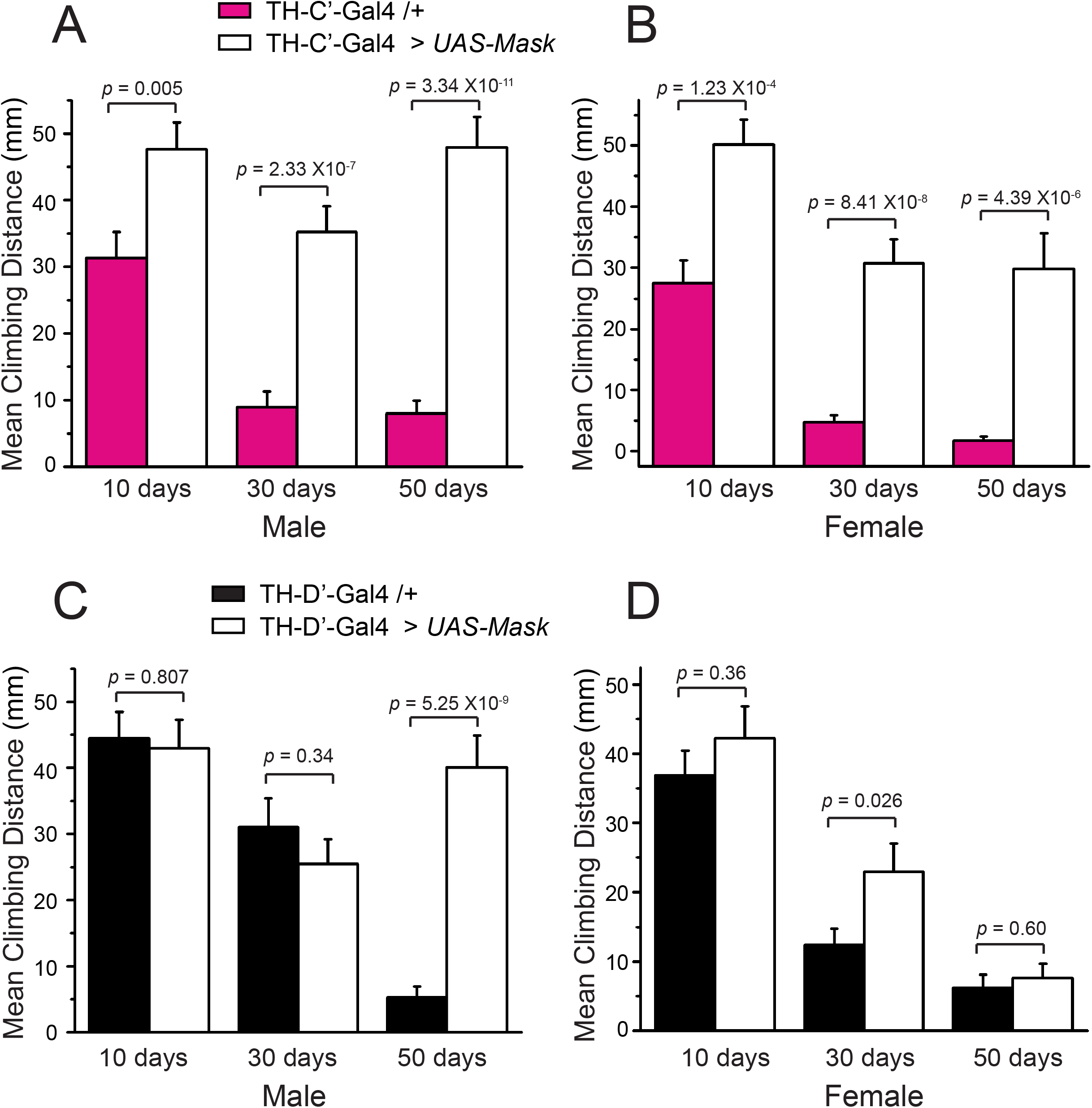
Overexpressing Mask in the TH-C’ and TH-D’ DANs improves locomotor in aged flies. Mean climb distance in Rapid Iterative Negative Geotaxis (RING) assay were measured to assess the locomotor in adult flies. Mean climb distance of a group of 15-20 A) male or B) female flies at ages of day-10, 30 or 50 are shown. Flies overexpressing Mask in the TH-C’ DANs show significantly faster climbing throughout the entire lifespan compared to the control flies. Mean climb distance of C) male and D) female control flies and flies overexpressing Mask in TH-D’ DA neurons were measured at ages of day-10, 30 or 50. Flies overexpressing Mask in the TH-D’ DA neurons are significantly more active at mid and old ages. (n=3 for all sample groups) **Note:** The RING Assay performed on 10- and 30-day old flies used 3 seconds total duration, while RING Assays on 50-day old flies used 5 seconds duration.

**Figure 6.**
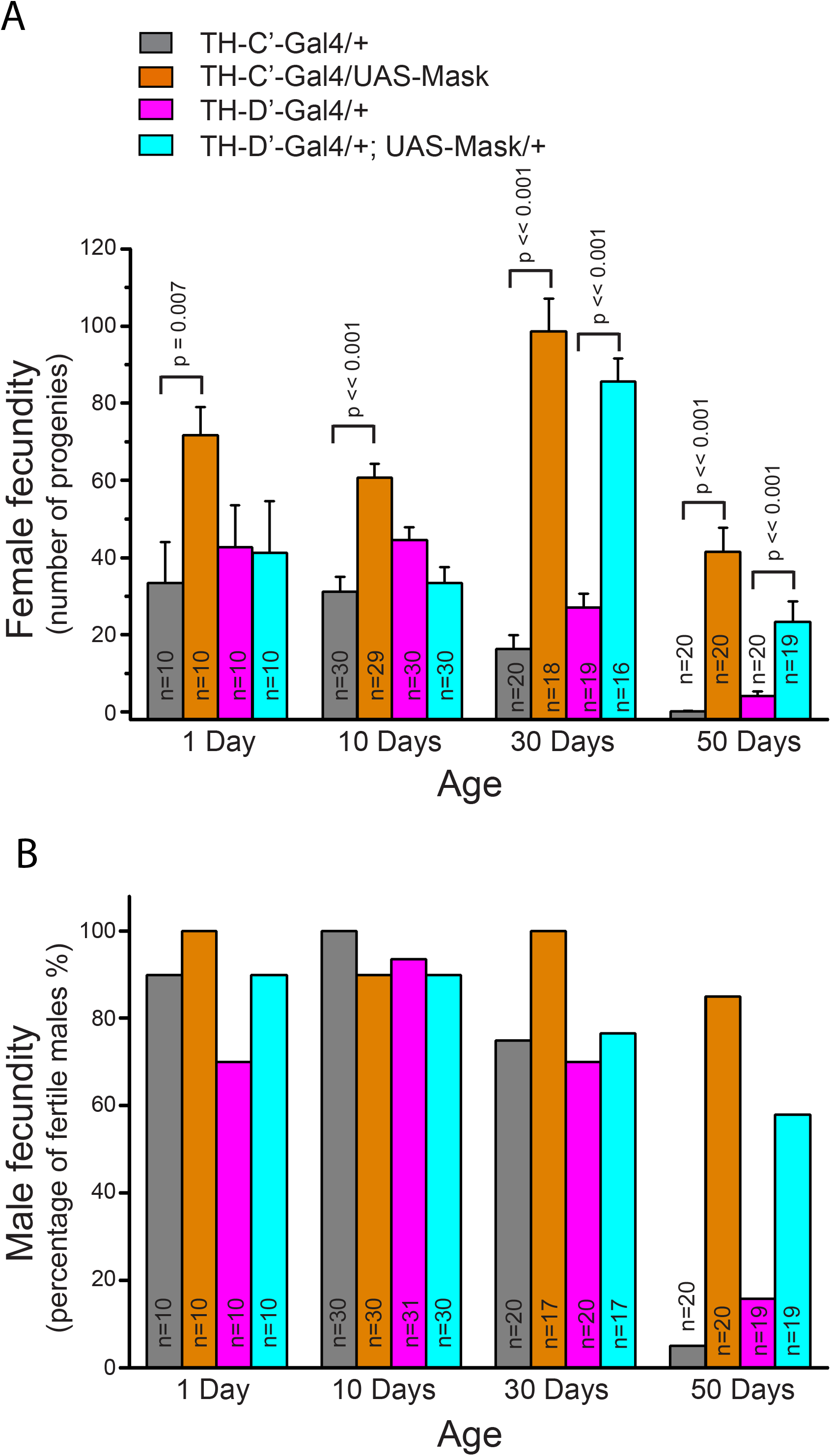
Mask-overexpressing Flies show sustained Fecundity at old ages. A) Numbers of progenies produced by female flies in a 3 day interval. The female flies that overexpress Mask in the TH-C’ or TH-D’ DANs produce significantly more progenies at 30- and 50-day old ages. B) Reproduction in male flies is measured as the percentage of male flies that are fertile. At age 50- day, majority of the control male flies lose the ability to produce progeny, while over 80% of male flies that overexpress Mask in the TH-C’ and 60% of those overexpress Mask in the TH-D’ DANs are still reproductively active.

### Tuning the microtubule (MT) stability in the DANs induces lifespan extension in flies

In a parallel study, we discovered that Mask promotes microtubule (MT) dynamics in fly larval motor neurons and body wall muscles^41^. I next tested whether altered MT dynamics in the Mask-expressing DANs is the key mediator for lifespan extension. Mask is a large protein (4001 amino acids) with many functional domains. As highlighted in Figure 7, Mask/ANKHD1 bears two Ankyrin Repeats clusters, one Nuclear Export Signal (NES) and one Nuclear Localization Signal (NLS), and a C-terminal KH domain. Ankyrin repeats mediate protein-protein interactions in eukaryotic cells^66^. The KH domain is a conserved motif that binds RNA or single-stranded DNA (ssDNA) and is found in proteins that are associated with transcription, splicing, translation and regulation of mRNA stability^67,68^. In order to better understand the molecular functions of Mask, we have generated several mutant Mask transgenes (depicted in Fig. 7A) to determine the structure-function correlations. One mutant transgene contains only the N-terminal ankryin repeats domains (Mask-ANK); one lacks the N-terminal half of the protein and contains the NES, NLS and KH domains (Mask-KH-Only); and the third one carries point mutations in the conserved amino acid “GXXG” motif^69^ in the KH domain (from GRGG to GDDG, named Mask-KH-Mut). Our structure-function analysis performed in fly larval muscles and motor neurons revealed that the Ankyrn repeats domain is necessary and sufficient to mediate the MT-regulating function of Mask^41^, while the KH domain is required for the autophagy-promoting function (Fig. S12). Using the same set of mutant Mask transgenes, I also determined the structural requirements for Mask to induce lifespan extension. The results showed that, when expressed in the TH-C’ DANs, the Mask-ANK and the Mask-KH-Mut transgenes both have the same ability as the wild type Mask protein to drive lifespan extension (Fig. 7B). In contrast, the Mask-KH only protein exhibits a dominant negative effect: overexpressing this truncated protein in the TH-C’ neurons reduces lifespan (Fig. 7B). These results suggest that the N-terminal portion of Mask protein containing the ankyrin repeats is necessary and sufficient to drive lifespan extension, while the NES, NLS and the KH domains are dispensable in this context. This tight structure and function correlations between the ability of the Ankryn repeats domains to promote MT dynamics and to induce lifespan extension indicates that MT dynamics is one of the primary contributing factors for the DANs-mediated beneficial effects on aging and longevity.

**Figure 7.**
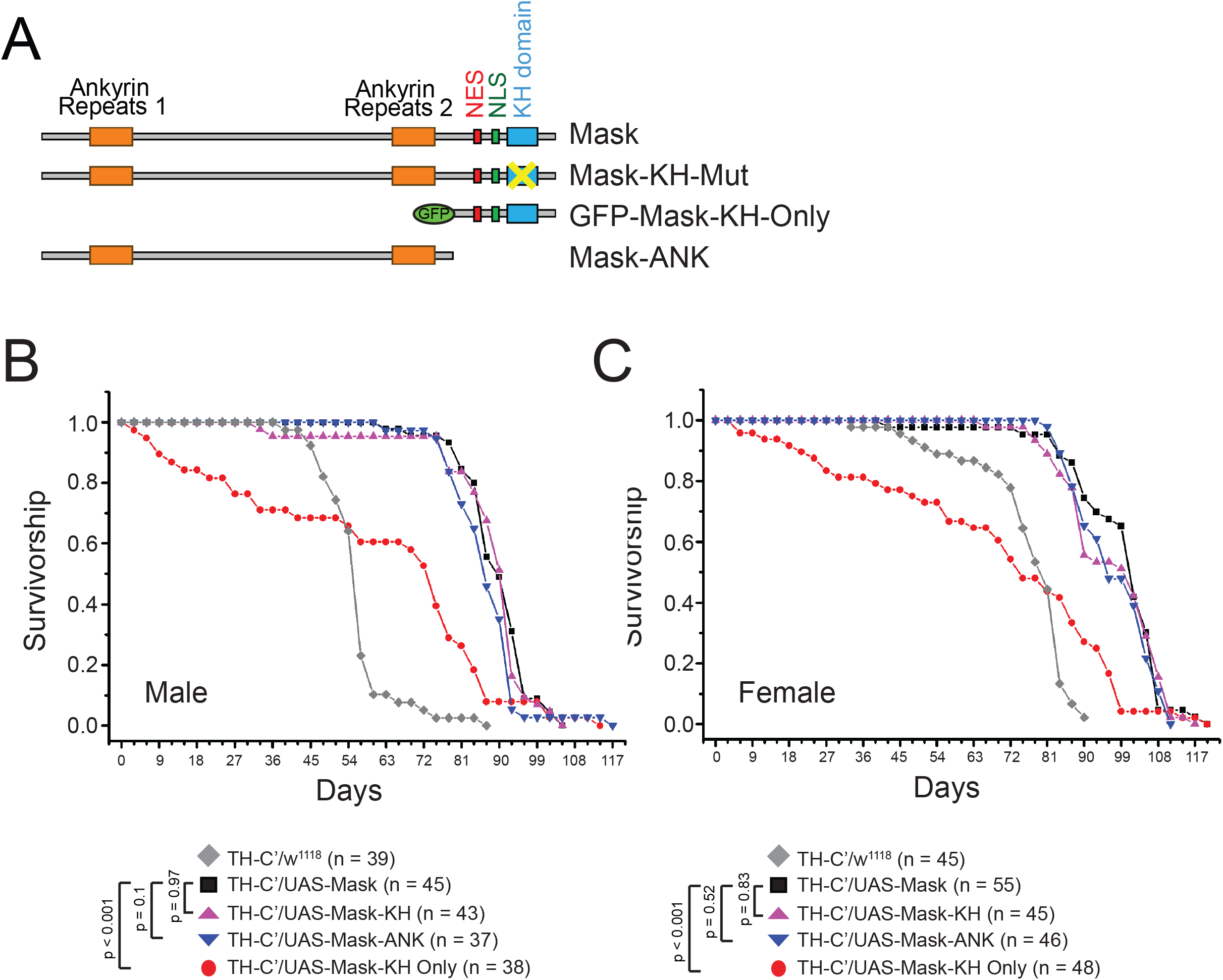
The Ankryn repeats domains in the Mask protein is sufficient to induce lifespan Extension. A) Schematic of domain structures of wild type and Mask mutant transgenes. B, C) Survivorship curves of male flies overexpressing different forms of Mask protein in the TH-C’ DANs. When overexpressed in the TH-C’ DANs, the Mask-ANK and Mask-KH-Mutant transgenes show similar strength to induce lifespan extension compared to the wild type Mask protein.

Based on this notion, one would expect that alternative interventions for MT dynamics would also be able to induce lifespan extension when applied to the same subsets of DANs. Activating the Kinesin heavy chain Unc-104^70,71^ and knocking down a component of the Dynein/Dynactin complex, p150^Glued72^, are two independent interventions that have been previously shown to increase MT dynamics^73^, and overexpressing Unc-104 in the nervous system has also been shown to moderately extend lifespan in worms^73^. In flies, the lifespan was extended by ~37% when Unc-104 is overexpressed in the TH-C’ DANs, and ~40% in the TH-D’ DANs (Fig. 8A). Reducing p150^Glued^ levels in the DANs exhibited similar but milder effects only when it’s applied to the TH-C’ DANs- the lifespan was extended by ~14%. These results together further demonstrate that increasing MT dynamics in the TH-C’ or TH-D’ DANs is sufficient to induce lifespan extension in flies.

**Figure 8.**
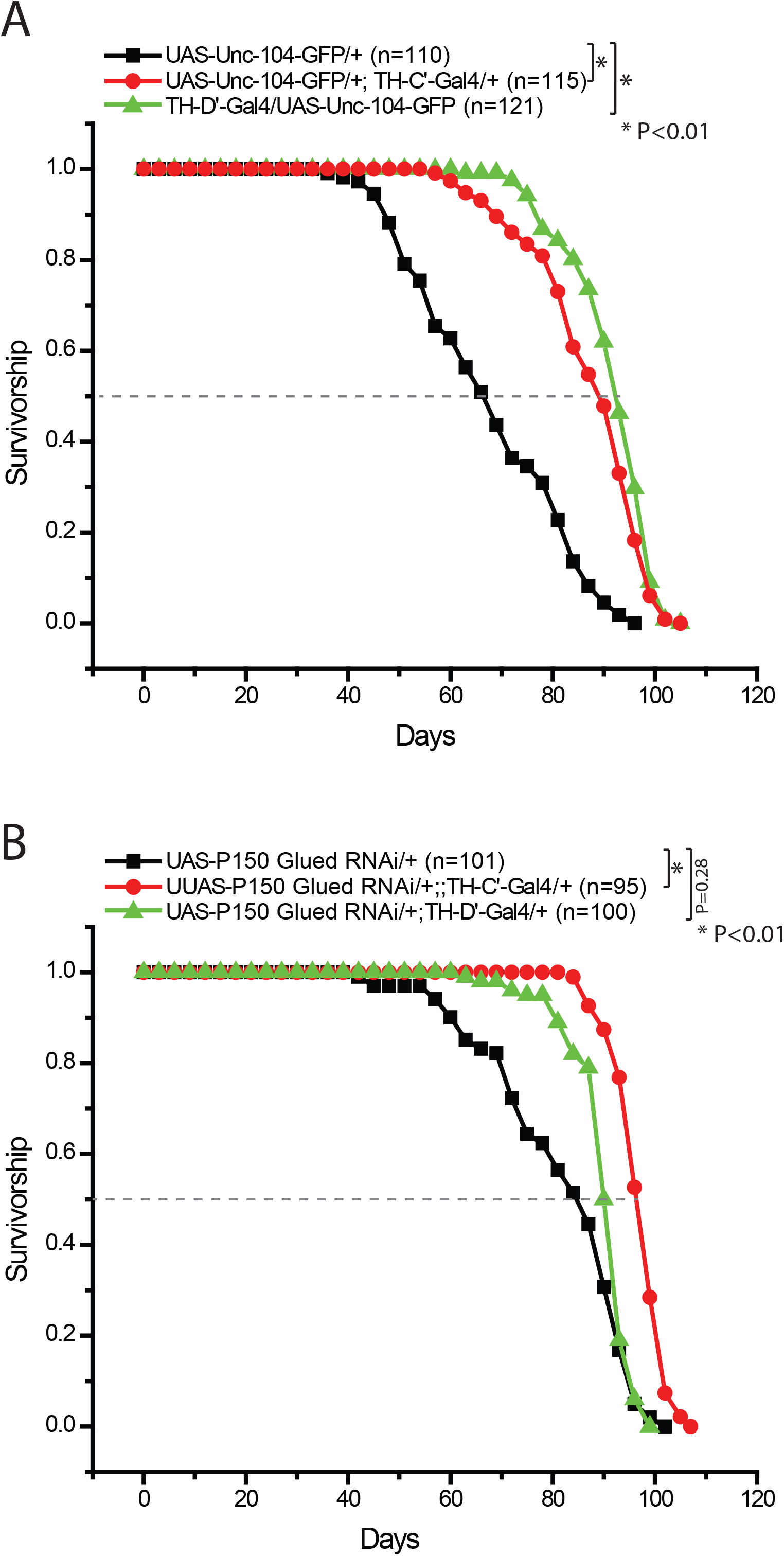
Overexpressing Unc-104 or Knocking down p150Glued in the TH-C’ or TH-D’ DANs extend lifespan in flies. Survivorship curves of female flies expressing UAS-GFP-Unc104 or a weak RNAi transgene for p150^Glued^ in the TH-C’ or TH-D’ DANs. A) Overexpressing the Unc-104-GFP transgene in the TH-C’ or TH-D’ neurons leads to extension of lifespan in flies. The median lifespan in the flies overexpressing Unc-104-GFP increased from ~67 days to ~92 days (~37% increase). In the flies overexpressing Unc-104-GFP in the TH-D’ neurons, the median lifespan increased ~40% (median lifespan ~94 days). B) Moderately knocking down p150^Glued^ in the TH-C’ neurons leads to extension of lifespan in flies, median lifespan increased from ~86 days to ~98 days (~13.95% increase). Same treatment in the TH-D’ neurons leads to an increase of the median lifespan to ~92 days (~6.97% increase).

## Discussion

In this study, I demonstrated that overexpressing the putative scaffolding protein Mask in two small groups of the DANs has a profound effect on longevity. Mask overexpression likely causes a gain-of-function effect that impacts the state of MT stability in these DANs. Conceivably, this effect would in turn induces a range of cellular changes that ultimately leads to altered neuronal outputs relayed to the peripheral tissues and resulting in improved aging and prolonged lifespan. These findings uncovered novel mechanism that modulates aging, but in order to fully understand this mechanism, many questions need to be answered. Among those, the most critical ones are: what are the peripheral tissues/organs that play a primary role to convey the impact on aging by specific DANs; and how and through what neuronal circuits do these DANs signal to the peripheral tissues?

The DANs are composed of a heterogeneous group of neurons with diverse morphology and functions^74^ that regulate a range of key brain activities. This work demonstrated that a specific small subsets of DANs are also able to regulate aging and longevity in flies. How might these small groups of DANs modulate systematic aging? One of the well-established DAN functions is the control of reward-feeding and energy balance at the whole animal level^24–26^. However, it is unclear whether DANs affect aging through the regulation of feeding and metabolism. Our analysis on the Mask-overexpressing long-lived flies indicated that the TH-C’ and TH-D’ DANs do not directly affect the amount of food consumption (Fig. S6). At the same time, the whole body TAG levels in these flies are elevated (Fig. S7), instead of being depleted as was indicated in the studies in worms that DAN-mediated regulation of lipid homeostasis underlies the related lifespan extension^18^. The TAG data, together with the sustained adult locomotor (Fig. 5), thus may reflect an extended health span in those long-lived Mask-overexpressing flies. Therefore, diet and metabolism are unlikely to be the primary underlying mechanism for the Mask-induced DAN-dependent lifespan extension. The Mask-induced DAN-dependent regulation on aging is also likely independent to disruptions in insulin production because the long-lived flies showed no consistent and relatable reduction of Dilp transcript levels in their brain (Fig S.8-11). This result is somewhat consistent with the finding that disruptions of the insulin pathway in worms enhances lifespan but has no effect on the age-dependent decline of dopamine levels^75^.

It is intriguing that the prolonged lifespan of the Mask-overexpressing flies is accompanied by a sustained fecundity in the long-lived flies (Fig. 6). Similar phenomena were also previously discovered by a number of other studies^62–65^. Moreover, reproductive female of eusocial insects show acquirable expansion of reproduction and lifespan. Together, these phenomenon challenge the long-noticed theory about the tradeoff between reproduction and longevity, and suggest that a mechanism capable of extend both reproduction and longevity may exist as a common strategy for animals to actively intervene aging process in order to cope with their reproductive demand. DANs, a key modulator for brain functions, may be a key element of a mechanism serves to coordinate the sustainment and fulfillment of reproduction in animals. Future investigation that further delineate the roles of DANs in regulating aging and reproduction can provide insight into the molecular and cellular basis for such a mechanism.

Mask and its mammalian homologues have been shown to play diverse functions in cells including signal transduction^35,76^, transcription^36,37^, autophagy^40^ and vesiculation^77^. Our most recent study demonstrated that Mask is also a potent regulator of MT stability (in revision). Which of these functions contribute to Mask-induced lifespan extension? Our structure-function analysis of the functional domains of Mask protein and our analysis of Mask-dependent manipulation of MT stability (Fig. 7) provided strong evidence suggesting that Mask’s ability to promote MT dynamics is a primary contributor for the DAN-mediated lifespan extension. This notion is further sported by the similar ability of Unc-104 overexpression or p150^Glued^ knockdown in the DANs to extend lifespan. It’s also worth noting that activation of Gα_s_ was also shown to enhance MT dynamics^78^, suggesting that the ability of moderately activating the engineered Gα_s_ receptor DREADD-Ds in the DANs to extend lifespan may share similar cellular mechanism as Mask-overexpression. Furthermore, I performed additional analysis to access whether Mask’s actions in regulating autophagy or vesiculation contribute to its ability to induce the dopamine-dependent lifespan extension. First, the Mask transgene carrying the KH-domain mutation fails to enhance autophagy (Fig. S12) but is sufficient to induce lifespan extension. Overexpressing ATG-1, a potent inducer of autophagy, in all neurons in the adult flies can extend lifespan^79^; however, overexpressing Mask in all neurons in the adult flies failed to induce similar effects (Fig. S13). These data argue against contribution of enhanced autophagy to Mask-induced lifespan extension. Second, although Mask is able to perform the conserved function as its mammalian counterpart in vesiculation^77^, as was indicated by the ability of Mask to increase the size of the presynaptic Rab5-positive vesicle pools (Fig. S14) and vesicular recycling (measured by FM-143 dye loading assay, see Fig. S15) at the larval NMJs, this function does not seem to be sufficient to induce lifespan extension. Overexpressing the active form of Rab5, which has been shown to enhance synaptic vesicle fusion efficacy and firing strength^80^, in the DANs could only moderately extend lifespan in the male flies, with a level that is much less significant compared to Mask-induced lifespan extension (Fig. S16). Thus, although Mask can enhance endocytosis and vesicle recycling at the nerve terminal, promoting these events alone in the DANs is not sufficient to induce the same outcome in lifespan extension.

MT provides structural platform for many essential cellular functions. This is particularly important for neurons both during development and in the adulthood for maintaining their normal activities. The stability and the balanced dynamics of the MTs are subjective to tight regulations in response to neuronal activities and stresses. Both normal and diseased aging of the brain are tightly linked to the dysregulations of MT stability and dynamics in neurons^81^. Given the broad contributions of the MT network to the integrity of neuronal structure and function, it is conceivable that altering the stability and enhancing the dynamics of the MT network in neurons could induce a wide range of cellular changes. These cellular changes could collectively alter neuronal behaviors and their circuitry outputs, improve their cellular homeostasis and mitigate the adverse effects occurred over age. The DANs, with their peculiar morphology, activity, metabolism and molecular signature, may be more susceptible to fine tunes of MT stability and dynamics.

## Method and Material

### Drosophila strains and genetics

Flies were maintained at 25°C on standard food containing agar (0.75% w/v), cornmeal (5% w/v), molasses (4.15% v/v), and yeast (1.575% w/v) at 25 °C in a 12:12 light:dark cycle. The following strains were used in this study: UAS-Mask^39^, TH-C’-Gal4^46^, TH-D’-Gal4^46^ (from Dr. Mark Wu), 0273-Gal4 (from Dr. Thomas R. Clandinin), UAS-DREADD-D_s_^47^ (from Charles Nichols), Ddc-Gal4 (Bloomington Stock #7010); UAS-unc104-GFP (Bloomington Stock #24787); UAS-p150^Glued^ RNAi (Bloomington Stock #24761). Control flies used in the study are *w^1118^* from BestGene Inc. (Chino Hills, CA, USA) in which the UAS-Mask line was generated^39^ and Iso^31^ (from Dr. Mark Wu) in which TH-C’-Gal4 and TH-D’-Gal4 flies were generated^46^.

### Fly Longevity Assay

The procedures are adapted from previously described protocol^82^. 15-20 male or female flies were separated into one vial, and a total of 4-6 vials of flies were used for each genotype for longevity measurement. The flies are maintained at 25°C on standard food or food containing drugs and are transferred into fresh vials every 3 days.

### DREADD Activation

10mM Clozapine N-oxide (CNO) (Tocris 4936) stock solution was prepared in DMSO. Food containing control (DMSO only) or 1uM CNO was prepared freshly every three days before the flies were transferred to new vials during the longevity recording experiments.

### Fly Fecundity Assay

Newly eclosed individual virgin male or female flies (day1) were crossed with 2-3 *w^1118^* virgin female or male flies respectively and kept together for 3 days. The total progeny produced within the 3-day intervals were counted after they eclose as adult flies. Sexually exposed and active male or female flies reared together with *w^1118^* female or male were aged till day 10, 30 and 50. Each female fly was then separated into a single vial and allow to lay eggs for 3 days. The total progeny produced within the 3-day intervals were counted after they eclose. The aged male flies were individually mated with three 5-7 day old *w^1118^* virgin females for 3 days, and fecundity was measured as the percentage of males that can successfully produce progenies.

### Statistical analysis

Data analysis for longevity: The Kaplan-Meier estimator was used to analyze the data and the survivorship curves were generated in Origin. The log-rank test was performed with Evan’s A/B tools.

Each sample was compared with other samples in the group (more than two) using one-way ANOVA, or with the other sample in a group of two using a t-test. The graphs were generated in Origin (Origin Lab, Northampton, MA).

## Supporting information

Tian_DAN-Mask_Longevity.PDF

## Acknowledgments

I greatly appreciate the help of Chunlai Wu for technical support and critical comments on the manuscript. I thank Hui Mao, Sarah Yanofski and Daniel Martinez for their technical support. I am especially grateful to Dr. Mark Wu for providing many helpful reagents. I also thank Dr. Charles Nichols and Dr. Thomas R. Clandinin for fly stocks, as well as the Bloomington Stock Center for other fly stocks and Developmental Studies Hybridoma Bank, created by the NICHD of the NIH and maintained at The University of Iowa, Department of Biology, Iowa City, IA 52242, for antibodies. I also thank Liz McGehee for editorial assistance. This work is supported by a start-up fund from LSUHSC to X.T.

## Author contributions

X.T. is responsible for experimental design, data collection and analysis. X.T. prepared the manuscript.

## Conflict of Interest Statement

The author certifies that she has NO affiliations with or involvement in any organization or entity with any financial interest, or non-financial interest in the subject matter or materials discussed in this manuscript.

## Data Availability Statement

The data that support the findings of this study are openly available in BioRxiv at https://www.biorxiv.org/content/10.1101/2020.06.15.153056v2

## Notes

### Competing Interest Statement

The authors have declared no competing interest.

### Summary of Updates

Reorganization of the data and manuscript and some editorial updates are included in the revised manuscript.

## References

1. Pasyukova E., Symonenko A., Roshina N., Trostnikov M., Veselkina E., Rybina O. Neuronal Genes and Developmental Neuronal Pathways in Drosophila Life Span Control. in Life Extension-lessons from Drosophila 3–37 (Springer, Cham, US, 2015).

2. Bishop, N. A. & Guarente, L. Two neurons mediate diet-restriction-induced longevity in C. elegans. Nature 447, 545–549 (2007). PubMed PMID: 17538612

3. Apfeld, J. & Kenyon, C. Regulation of lifespan by sensory perception in Caenorhabditis elegans. Nature 402, 804–809 (1999). PubMed PMID: 10617200

4. Alcedo, J. & Kenyon, C. Regulation of C. elegans longevity by specific gustatory and olfactory neurons. Neuron 41, 45–55 (2004). PubMed PMID: 14715134

5. Lee, S. J. & Kenyon, C. Regulation of the longevity response to temperature by thermosensory neurons in Caenorhabditis elegans. Curr Biol 19, 715–722 (2009). PubMed PMID: 19375320

6. Zhang, B., Gong, J., Zhang, W., Xiao, R., Liu, J. & Xu, X. Z. S. Brain-gut communications via distinct neuroendocrine signals bidirectionally regulate longevity in C. elegans. Genes Dev 32, 258–270 (2018). PubMed PMID: 29491136

7. Chen, Y. C., Chen, H. J., Tseng, W. C., Hsu, J. M., Huang, T. T., Chen, C. H. & Pan, C. L. A C. elegans Thermosensory Circuit Regulates Longevity through crh-1/CREB-Dependent flp-6 Neuropeptide Signaling. Dev Cell 39, 209–223 (2016). PubMed PMID: 27720609

8. Libert, S., Zwiener, J., Chu, X., Vanvoorhies, W., Roman, G. & Pletcher, S. D. Regulation of Drosophila life span by olfaction and food-derived odors. Science 315, 1133–1137 (2007). PubMed PMID: 17272684

9. Waterson, M. J., Chung, B. Y., Harvanek, Z. M., Ostojic, I., Alcedo, J. & Pletcher, S. D. Water sensor ppk28 modulates Drosophila lifespan and physiology through AKH signaling. Proc Natl Acad Sci U S A 111, 8137–8142 (2014). PubMed PMID: 24821805

10. Gendron, C. M., Kuo, T. H., Harvanek, Z. M., Chung, B. Y., Yew, J. Y., Dierick, H. A. & Pletcher, S. D. Drosophila life span and physiology are modulated by sexual perception and reward. Science 343, 544–548 (2014). PubMed PMID: 24292624

11. Broughton, S. J., Piper, M. D., Ikeya, T., Bass, T. M., Jacobson, J., Driege, Y., Martinez, P., Hafen, E., Withers, D. J., Leevers, S. J. & Partridge, L. Longer lifespan, altered metabolism, and stress resistance in Drosophila from ablation of cells making insulin-like ligands. Proc Natl Acad Sci U S A 102, 3105–3110 (2005). PubMed PMID: 15708981

12. Broughton, S. J., Slack, C., Alic, N., Metaxakis, A., Bass, T. M., Driege, Y. & Partridge, L. DILP-producing median neurosecretory cells in the Drosophila brain mediate the response of lifespan to nutrition. Aging Cell 9, 336–346 (2010). PubMed PMID: 20156206

13. Zhang, G., Li, J., Purkayastha, S., Tang, Y., Zhang, H., Yin, Y., Li, B., Liu, G. & Cai, D. Hypothalamic programming of systemic ageing involving IKK-beta, NF-kappaB and GnRH. Nature 497, 211–216 (2013). PubMed PMID: 23636330

14. Zhang, Y., Kim, M. S., Jia, B., Yan, J., Zuniga-Hertz, J. P., Han, C. & Cai, D. Hypothalamic stem cells control ageing speed partly through exosomal miRNAs. Nature 548, 52–57 (2017). PubMed PMID: 28746310

15. Satoh, A. & Imai, S. Systemic regulation of mammalian ageing and longevity by brain sirtuins. Nat Commun 5, 4211 (2014). PubMed PMID: 24967620

16. Bolukbasi, E., Woodling, N. S., Ivanov, D. K., Adcott, J., Foley, A., Rajasingam, A., Gittings, L. M., Aleyakpo, B., Niccoli, T., Thornton, J. M. & Partridge, L. Cell type-specific modulation of healthspan by Forkhead family transcription factors in the nervous system. Proc Natl Acad Sci U S A 118(2021). PubMed PMID: 33593901

17. Zhang, Q., Wu, X., Chen, P., Liu, L., Xin, N., Tian, Y. & Dillin, A. The Mitochondrial Unfolded Protein Response Is Mediated Cell-Non-autonomously by Retromer-Dependent Wnt Signaling. Cell 174, 870–883.e817 (2018). PubMed PMID: 30057120

18. Higuchi-Sanabria, R., Durieux, J., Kelet, N., Homentcovschi, S., de Los Rios Rogers, M., Monshietehadi, S., Garcia, G., Dallarda, S., Daniele, J. R., Ramachandran, V., Sahay, A., Tronnes, S. U., Joe, L. & Dillin, A. Divergent Nodes of Non-autonomous UPR(ER) Signaling through Serotonergic and Dopaminergic Neurons. Cell Rep 33, 108489 (2020). PubMed PMID: 33296657

19. Lyu, Y., Weaver, K. J., Shaukat, H. A., Plumoff, M. L., Tjilos, M., Promislow, D. E. & Pletcher, S. D. Drosophila serotonin 2A receptor signaling coordinates central metabolic processes to modulate aging in response to nutrient choice. Elife 10(2021). PubMed PMID: 33463526

20. Chakraborty, T. S., Gendron, C. M., Lyu, Y., Munneke, A. S., DeMarco, M. N., Hoisington, Z. W. & Pletcher, S. D. Sensory perception of dead conspecifics induces aversive cues and modulates lifespan through serotonin in Drosophila. Nat Commun 10, 2365 (2019). PubMed PMID: 31147540

21. Ro, J., Pak, G., Malec, P. A., Lyu, Y., Allison, D. B., Kennedy, R. T. & Pletcher, S. D. Serotonin signaling mediates protein valuation and aging. Elife 5(2016). PubMed PMID: 27572262

22. Schultz, W. Multiple dopamine functions at different time courses. Annu Rev Neurosci 30, 259–288 (2007). PubMed PMID: 17600522

23. Arias-Carrión, O., Stamelou, M., Murillo-Rodríguez, E., Menéndez-González, M. & Pöppel, E. Dopaminergic reward system: a short integrative review. Int Arch Med 3, 24 (2010). PubMed PMID: 20925949

24. Murray, S., Tulloch, A., Gold, M. S. & Avena, N. M. Hormonal and neural mechanisms of food reward, eating behaviour and obesity. Nat Rev Endocrinol 10, 540–552 (2014). PubMed PMID: 24958311

25. Narayanan, N. S., Guarnieri, D. J. & DiLeone, R. J. Metabolic hormones, dopamine circuits, and feeding. Front Neuroendocrinol 31, 104–112 (2010). PubMed PMID: 19836414

26. Doan, K. V., Kinyua, A. W., Yang, D. J., Ko, C. M., Moh, S. H., Shong, K. E., Kim, H., Park, S. K., Kim, D. H., Kim, I., Paik, J. H., DePinho, R. A., Yoon, S. G., Kim, I. Y., Seong, J. K., Choi, Y. H. & Kim, K. W. FoxO1 in dopaminergic neurons regulates energy homeostasis and targets tyrosine hydroxylase. Nat Commun 7, 12733 (2016). PubMed PMID: 27681312

27. Cotzias, G. C., Miller, S. T., Nicholson, A. R., Jr., Maston, W. H. & Tang, L. C. Prolongation of the life-span in mice adapted to large amounts of L-dopa. Proc Natl Acad Sci U S A 71, 2466–2469 (1974). PubMed PMID: 4526220

28. Knoll, J. Sexual performance and longevity. Exp Gerontol 32, 539–552 (1997). PubMed PMID: 9315455

29. De Luca, M., Roshina, N. V., Geiger-Thornsberry, G. L., Lyman, R. F., Pasyukova, E. G. & Mackay, T. F. Dopa decarboxylase (Ddc) affects variation in Drosophila longevity. Nat Genet 34, 429–433 (2003). PubMed PMID: 12881721

30. Riemensperger, T., Isabel, G., Coulom, H., Neuser, K., Seugnet, L., Kume, K., Iche-Torres, M., Cassar, M., Strauss, R., Preat, T., Hirsh, J. & Birman, S. Behavioral consequences of dopamine deficiency in the Drosophila central nervous system. Proc Natl Acad Sci U S A 108, 834–839 (2011). PubMed PMID: 21187381

31. Murakami, H. & Murakami, S. Serotonin receptors antagonistically modulate Caenorhabditis elegans longevity. Aging Cell 6, 483–488 (2007). PubMed PMID: 17559503

32. Savica, R., Grossardt, B. R., Bower, J. H., Ahlskog, J. E., Boeve, B. F., Graff-Radford, J., Rocca, W. A. & Mielke, M. M. Survival and Causes of Death Among People With Clinically Diagnosed Synucleinopathies With Parkinsonism: A Population-Based Study. JAMA Neurol 74, 839–846 (2017). PubMed PMID: 28505261

33. Yamamoto, Branden J. Stansley & Bryan K. L-Dopa and Brain Serotonin System Dysfunction. Toxics 3, 75–88 (2015). PubMed PMID:

34. Lees, A. J. Comparison of therapeutic effects and mortality data of levodopa and levodopa combined with selegiline in patients with early, mild Parkinson’s disease. Parkinson’s Disease Research Group of the United Kingdom. Bmj 311, 1602–1607 (1995). PubMed PMID: 8555803

35. Smith, R. K., Carroll, P. M., Allard, J. D. & Simon, M. A. MASK, a large ankyrin repeat and KH domain-containing protein involved in Drosophila receptor tyrosine kinase signaling. Development 129, 71–82 (2002). PubMed PMID: 11782402

36. Sansores-Garcia, L., Atkins, M., Moya, I. M., Shahmoradgoli, M., Tao, C., Mills, G. B. & Halder, G. Mask is required for the activity of the Hippo pathway effector Yki/YAP. Curr Biol 23, 229–235 (2013). PubMed PMID: 23333314

37. Sidor, C. M., Brain, R. & Thompson, B. J. Mask proteins are cofactors of Yorkie/YAP in the Hippo pathway. Curr Biol 23, 223–228 (2013). PubMed PMID: 23333315

38. Fisher, K. H., Fragiadaki, M., Pugazhendhi, D., Bausek, N., Arredondo, M. A., Thomas, S. J., Brown, S. & Zeidler, M. P. A genome-wide RNAi screen identifies MASK as a positive regulator of cytokine receptor stability. J Cell Sci (2018). PubMed PMID: 29848658

39. Zhu, M., Li, X., Tian, X. & Wu, C. Mask loss-of-function rescues mitochondrial impairment and muscle degeneration of Drosophila pink1 and parkin mutants. Hum Mol Genet 24, 3272–3285 (2015). PubMed PMID: 25743185

40. Zhu, M., Zhang, S., Tian, X. & Wu, C. Mask mitigates MAPT- and FUS-induced degeneration by enhancing autophagy through lysosomal acidification. Autophagy 13, 1924–1938 (2017). PubMed PMID: 28806139

41. Martinez, Daniel, Zhu, Mingwei, Guidry, Jessie J., Majeste, Niles, Mao, Hui, Yanofsky, Sarah, Tian, Xiaolin & Wu, Chunlai. Mask, the *Drosophila* Ankyrin Repeat and KH domain-containing protein, regulates microtubule dynamics. bioRxiv, 2020.2004.2022.056051 (2021). PubMed PMID:

42. Zhu, M., Zhang, S., Tian, X. & Wu, C. Mask mitigates MAPT- and FUS-induced degeneration by enhancing autophagy through lysosomal acidification. Autophagy, 1–15 (2017). PubMed PMID: 28806139

43. Mao, Z. & Davis, R. L. Eight different types of dopaminergic neurons innervate the Drosophila mushroom body neuropil: anatomical and physiological heterogeneity. Front Neural Circuits 3, 5 (2009). PubMed PMID: 19597562

44. Burke, C. J., Huetteroth, W., Owald, D., Perisse, E., Krashes, M. J., Das, G., Gohl, D., Silies, M., Certel, S. & Waddell, S. Layered reward signalling through octopamine and dopamine in Drosophila. Nature 492, 433–437 (2012). PubMed PMID: 23103875

45. Galili, D. S., Dylla, K. V., Ludke, A., Friedrich, A. B., Yamagata, N., Wong, J. Y., Ho, C. H., Szyszka, P. & Tanimoto, H. Converging circuits mediate temperature and shock aversive olfactory conditioning in Drosophila. Curr Biol 24, 1712–1722 (2014). PubMed PMID: 25042591

46. Liu, Q., Liu, S., Kodama, L., Driscoll, M. R. & Wu, M. N. Two dopaminergic neurons signal to the dorsal fan-shaped body to promote wakefulness in Drosophila. Curr Biol 22, 2114–2123 (2012). PubMed PMID: 23022067

47. Becnel, J., Johnson, O., Majeed, Z. R., Tran, V., Yu, B., Roth, B. L., Cooper, R. L., Kerut, E. K. & Nichols, C. D. DREADDs in Drosophila: a pharmacogenetic approach for controlling behavior, neuronal signaling, and physiology in the fly. Cell Rep 4, 1049–1059 (2013). PubMed PMID: 24012754

48. Granger, A. J., Wallace, M. L. & Sabatini, B. L. Multi-transmitter neurons in the mammalian central nervous system. Curr Opin Neurobiol 45, 85–91 (2017). PubMed PMID: 28500992

49. Aso, Y., Ray, R. P., Long, X., Bushey, D., Cichewicz, K., Ngo, T. T., Sharp, B., Christoforou, C., Hu, A., Lemire, A. L., Tillberg, P., Hirsh, J., Litwin-Kumar, A. & Rubin, G. M. Nitric oxide acts as a cotransmitter in a subset of dopaminergic neurons to diversify memory dynamics. Elife 8(2019). PubMed PMID: 31724947

50. Aguilar, J. I., Dunn, M., Mingote, S., Karam, C. S., Farino, Z. J., Sonders, M. S., Choi, S. J., Grygoruk, A., Zhang, Y., Cela, C., Choi, B. J., Flores, J., Freyberg, R. J., McCabe, B. D., Mosharov, E. V., Krantz, D. E., Javitch, J. A., Sulzer, D., Sames, D., Rayport, S. & Freyberg, Z. Neuronal Depolarization Drives Increased Dopamine Synaptic Vesicle Loading via VGLUT. Neuron 95, 1074–1088.e1077 (2017). PubMed PMID: 28823729

51. Kayser, M. S., Yue, Z. & Sehgal, A. A critical period of sleep for development of courtship circuitry and behavior in Drosophila. Science 344, 269–274 (2014). PubMed PMID: 24744368

52. Walters, M. R., Dutertre, M. & Smith, C. L. SKF-82958 is a subtype-selective estrogen receptor-alpha (ERalpha) agonist that induces functional interactions between ERalpha and AP-1. J Biol Chem 277, 1669–1679 (2002). PubMed PMID: 11700319

53. Wiemerslage, L., Schultz, B. J., Ganguly, A. & Lee, D. Selective degeneration of dopaminergic neurons by MPP(+) and its rescue by D2 autoreceptors in Drosophila primary culture. J Neurochem 126, 529–540 (2013). PubMed PMID: 23452092

54. Liu, Q., Tabuchi, M., Liu, S., Kodama, L., Horiuchi, W., Daniels, J., Chiu, L., Baldoni, D. & Wu, M. N. Branch-specific plasticity of a bifunctional dopamine circuit encodes protein hunger. Science 356, 534–539 (2017). PubMed PMID: 28473588

55. Tsao, C. H., Chen, C. C., Lin, C. H., Yang, H. Y. & Lin, S. Drosophila mushroom bodies integrate hunger and satiety signals to control innate food-seeking behavior. Elife 7(2018). PubMed PMID: 29547121

56. Landayan, D. & Wolf, F. W. Shared neurocircuitry underlying feeding and drugs of abuse in Drosophila. Biomed J 38, 496–509 (2015). PubMed PMID: 27013449

57. Marella, S., Mann, K. & Scott, K. Dopaminergic modulation of sucrose acceptance behavior in Drosophila. Neuron 73, 941–950 (2012). PubMed PMID: 22405204

58. Diegelmann, S., Jansen, A., Jois, S., Kastenholz, K., Velo Escarcena, L., Strudthoff, N. & Scholz, H. The CApillary FEeder Assay Measures Food Intake in Drosophila melanogaster. J Vis Exp (2017). PubMed PMID: 28362419

59. Nichols, C. D., Becnel, J. & Pandey, U. B. Methods to assay Drosophila behavior. J Vis Exp (2012). PubMed PMID: 22433384

60. Gargano, J. W., Martin, I., Bhandari, P. & Grotewiel, M. S. Rapid iterative negative geotaxis (RING): a new method for assessing age-related locomotor decline in Drosophila. Exp Gerontol 40, 386–395 (2005). PubMed PMID: 15919590

61. Yuan, R., Meng, Q., Nautiyal, J., Flurkey, K., Tsaih, S. W., Krier, R., Parker, M. G., Harrison, D. E. & Paigen, B. Genetic coregulation of age of female sexual maturation and lifespan through circulating IGF1 among inbred mouse strains. Proc Natl Acad Sci U S A 109, 8224–8229 (2012). PubMed PMID: 22566614

62. Simon, A. F., Shih, C., Mack, A. & Benzer, S. Steroid control of longevity in Drosophila melanogaster. Science 299, 1407–1410 (2003). PubMed PMID: 12610309

63. Hwangbo, D. S., Gershman, B., Tu, M. P., Palmer, M. & Tatar, M. Drosophila dFOXO controls lifespan and regulates insulin signalling in brain and fat body. Nature 429, 562–566 (2004). PubMed PMID: 15175753

64. Marden, J. H., Rogina, B., Montooth, K. L. & Helfand, S. L. Conditional tradeoffs between aging and organismal performance of Indy long-lived mutant flies. Proc Natl Acad Sci U S A 100, 3369–3373 (2003). PubMed PMID: 12626742

65. Grandison, R. C., Piper, M. D. & Partridge, L. Amino-acid imbalance explains extension of lifespan by dietary restriction in Drosophila. Nature 462, 1061–1064 (2009). PubMed PMID: 19956092

66. Mosavi, L. K., Cammett, T. J., Desrosiers, D. C. & Peng, Z. Y. The ankyrin repeat as molecular architecture for protein recognition. Protein Sci 13, 1435–1448 (2004). PubMed PMID: 15152081

67. Garcia-Mayoral, M. F., Hollingworth, D., Masino, L., Diaz-Moreno, I., Kelly, G., Gherzi, R., Chou, C. F., Chen, C. Y. & Ramos, A. The structure of the C-terminal KH domains of KSRP reveals a noncanonical motif important for mRNA degradation. Structure 15, 485–498 (2007). PubMed PMID: 17437720

68. Grishin, N. V. KH domain: one motif, two folds. Nucleic Acids Res 29, 638–643 (2001). PubMed PMID: 11160884

69. Hollingworth, D., Candel, A. M., Nicastro, G., Martin, S. R., Briata, P., Gherzi, R. & Ramos, A. KH domains with impaired nucleic acid binding as a tool for functional analysis. Nucleic Acids Res 40, 6873–6886 (2012). PubMed PMID: 22547390

70. Randall, T. S., Yip, Y. Y., Wallock-Richards, D. J., Pfisterer, K., Sanger, A., Ficek, W., Steiner, R. A., Beavil, A. J., Parsons, M. & Dodding, M. P. A small-molecule activator of kinesin-1 drives remodeling of the microtubule network. Proc Natl Acad Sci U S A 114, 13738–13743 (2017). PubMed PMID: 29229862

71. Straube, A., Hause, G., Fink, G. & Steinberg, G. Conventional kinesin mediates microtubule-microtubule interactions in vivo. Mol Biol Cell 17, 907–916 (2006). PubMed PMID: 16339079

72. Lazarus, J. E., Moughamian, A. J., Tokito, M. K. & Holzbaur, E. L. Dynactin subunit p150(Glued) is a neuron-specific anti-catastrophe factor. PLoS Biol 11, e1001611 (2013). PubMed PMID: 23874158

73. Li, L. B., Lei, H., Arey, R. N., Li, P., Liu, J., Murphy, C. T., Xu, X. Z. & Shen, K. The Neuronal Kinesin UNC-104/KIF1A Is a Key Regulator of Synaptic Aging and Insulin Signaling-Regulated Memory. Curr Biol 26, 605–615 (2016). PubMed PMID: 26877087

74. Vogt Weisenhorn, D. M., Giesert, F. & Wurst, W. Diversity matters - heterogeneity of dopaminergic neurons in the ventral mesencephalon and its relation to Parkinson’s Disease. J Neurochem 139 Suppl 1, 8–26 (2016). PubMed PMID: 27206718

75. Yin, J. A., Liu, X. J., Yuan, J., Jiang, J. & Cai, S. Q. Longevity manipulations differentially affect serotonin/dopamine level and behavioral deterioration in aging Caenorhabditis elegans. J Neurosci 34, 3947–3958 (2014). PubMed PMID: 24623772

76. Fisher, K. H., Fragiadaki, M., Pugazhendhi, D., Bausek, N., Arredondo, M. A., Thomas, S. J., Brown, S. & Zeidler, M. P. A genome-wide RNAi screen identifies MASK as a positive regulator of cytokine receptor stability. J Cell Sci 131(2018). PubMed PMID: 29848658

77. Kitamata, M., Hanawa-Suetsugu, K., Maruyama, K. & Suetsugu, S. Membrane-Deformation Ability of ANKHD1 Is Involved in the Early Endosome Enlargement. iScience 17, 101–118 (2019). PubMed PMID: 31255983

78. Sierra-Fonseca, Sukla Roychowdhury and Jorge A. Heterotrimeric G Proteins and the Regulation of Microtubule Assembly. in Cytoskeleton Structure, Dynamics, Function and Disease (ed. Jimenez-Lopez, J.C.).

79. Ulgherait, M., Rana, A., Rera, M., Graniel, J. & Walker, D. W. AMPK modulates tissue and organismal aging in a non-cell-autonomous manner. Cell Rep 8, 1767–1780 (2014). PubMed PMID: 25199830

80. Wucherpfennig, T., Wilsch-Brauninger, M. & Gonzalez-Gaitan, M. Role of Drosophila Rab5 during endosomal trafficking at the synapse and evoked neurotransmitter release. J Cell Biol 161, 609–624 (2003). PubMed PMID: 12743108

81. Baas, P. W., Rao, A. N., Matamoros, A. J. & Leo, L. Stability properties of neuronal microtubules. Cytoskeleton (Hoboken) 73, 442–460 (2016). PubMed PMID: 26887570

82. Linford, N. J., Bilgir, C., Ro, J. & Pletcher, S. D. Measurement of lifespan in Drosophila melanogaster. J Vis Exp (2013). PubMed PMID: 23328955

