## Supplementary material for "Enhancing Mask Activity in Dopaminergic Neurons Extends Lifespan in Flies": Tian_DAN-Mask_Longevity.PDF

**Supplemental Information**

**SI Methods and Material**

*Fly stocks*

The following stocks were used: UAS-*mask* RNAi[1], elav-Gal4[2] (neuron specific), Repo-Gal4 (glial specific); DA-Gal4 (Ubiquitous), UAS-DREADD-Di (from Charles Nichols), UAS-MASK-KH-Mutant (this study). Stocks from the Bloomington stock center include UAS-Ple or UAS-TH (37539), UAS-Ddc (37540), UAS-tdTomato (64746), UAS-TrpA1 (26263), $ple^{4}$ (3279), $ddc^{27}$ (3190), $dat^{Z2\text{-}1744}$ (30867), $vmat^{SH0459}$ (29477), UAS-GFP-mCherry-ATG8 (37749), GS-elav (43642), UAS-ATG1 (51654), UAS-YFP-Rab5 (9775), UAS-YFP-Rab5-S43N (9771), UAS-YFP-Rab5-Q88L (9774).

*Drug Treatment*

Dopamine pathway activation: microwave melted pre-made standard fly food was blended with drugs or the same amount of water or their solvent. The final concentrations for the drugs were used in this study: 1mg/ml L-DOPA[3,4] (3mg/ml L-DOPA stock in water, 0.1% L-ascorbic acid to prevent oxidation of L-DOPA); 30uM Quinpirole[5] ( 30mM Quinpirole stock was prepared in water); and 75uM SKF-82958[6] (15mM SKF-82958 stock was prepared in water).

Gene Switch system: Flies were raised on RU-486 (50ug/ml) food for the entire lifespan, control group were kept on food added equal amount of ethanol (10mg/ml RU486 stock was dissolved in ethanol).

*Generation of UAS-Mask mutant Transgenes*

The full-length wild type *mask* cDNA[1] was used to generate mutant and truncated *mask* cDNA, including pUAST-Mask-KH-Mut, pUAST-GFP-Mask-KH-Only, pUAST-Mask-Ank. To generate the UAS-Mask-KH-Mut transgenes, a complementary primer pair containing the mutated coding sequence (GGA<u>GACGAT</u>GGA, mutations underlined) were used to facilitate the replacement of the amino acid sequence (AA3053-3056) from GRGG in the wild type Mask to GDDG in the Mask-KH-Mut transgene. We used the QuickChange II XL site-mutagenesis kit (from Agilent Technology, Santa Clara, CA) to substitute wild type sequence with the mutant sequence. The entire UAS-Mask-KH-Mut coding region was then sequenced to ensure no unintended mutations were introduced. To generate the UAS-Mask-ANK and UAS-Mask-KH-Only transgenes, the KpnI (8001) site was used to separate the Mask encoding region into an N- and a C-terminal fragment, each was then used to generate the pUAST-Mask-ANK and pUAST-Mask-KH-Only respectively. All transgenic fly lines were generated by BestGene Inc. (Chino Hills, CA, USA).

*Glucose and TAG measurement*

Five male or female flies were collected and snap frozen in liquid nitrogen. The flies were homogenized in 200µL lysis buffer containing 1XPBS + 0.05% Triton X-100, centrifuged at 1,000×g for 1 min, and the supernatant was used to measure the protein, glucose and triacylglyceride (TAG) concentrations. 5 µL homogenate was used for TAG measurement using the Infinity Trigly-cerides reagent (Thermo Scientific). The sample was added to warmed reagent, mixed briefly, incubated at room temperature for 5 minutes, and read at 520 nm. For glucose measurement, 10µL homogenate was used to mix with 150 µL warmed Infinity Glucose reagent (Thermo Scientific), incubate for 30 min at 37 °C, and read at 340nm. For protein measurement, 2μL homogenate was used to determine the protein concentration using DC detection kit (BioRad). All reactions were read with SpectraMAX 190 (Molecular Device).

*Fly larval Neuromuscular Junction FM1-43 Dye loading*

The 3rd instar wondering larvae were dissected in HL3 buffer that contains 0.43uM $Ca^{2+}$ and then incubated in Jan's high $K^+$ buffer (90mM $K^+$, 2mM $Ca^{2+}$) + 4uM FM1-43 dye (final concentration) for 1 minute. The FM1-43 loading was then stopped by rinsing the larvae in $Ca^{2+}$ free HL3 buffer rapidly twice and then slowly on a shaker for 5 minutes. The larvae were then fixed in 4% PFA (in 1X PBS) for 18 minutes, rapidly rinsed in 1X PBS twice and then incubated in HRP-Cy5 (1:1,000) for 15 minutes. The larvae then were rapidly rinsed with PBS twice and followed by a slow rinse by shaking for 5 minutes. After the rinses, the larvae were incubated in 70% glycerol (in 1X PBS) for 15 minutes and mounted for confocal imaging analysis.

*Immunocytochemistry*

Third instar larvae were dissected in ice-cold PBS and fixed in 4% PFA for 30 min. The fixed tissues were stained following standard procedures. Alexa-Fluor-647-conjugated goat anti-HRP at 1:1000 was used to highlight the neuronal processes.

*Confocal imaging and analysis*

Single-layer or z-stack confocal images were captured on a Nikon (Tokyo, Japan) C1 confocal microscope. Images shown in the same figure were acquired using the same gain from samples that had been simultaneously fixed and stained. For quantification of YFP and FM1-43 signals at the NMJs, z-stack confocal images of microtubule in larval muscle 6/7 in segment A2 were analyzed double-blinded, IMARIS software (Bitplane, Inc) was used to quantify average signal intensity at the nerve terminals.

*Fly in vivo assays*

The procedures of Capillary Feeder assay (CAFE) was adapted from established protocols[7].

*qRT_PCR*

20 adult fly heads were collected, rinse with PBS 3 times and then snap frozen in liquid nitrogen. The total RNA was extracted with Trizol reagent. To prepare cDNA, the RNA samples were first treated with DNase I (Invitrogen Cat#2020-08-31), and then used as template to synthesize the cDNA using the High Capacity cDNA Revers Transcription kit (Applied Biosystem, Cat#2020-05-31). The quantitative PCR was performed with the BioRad SYBR (Cat# 1725121) and CFX96 Real-time PCR system.

Primers used in qPCR: dilp2, dilp5, dilp5 dilp6. Primers are described previously[8];

Primers for internal control: (reference: DOI: 10.1038/ncomms10368) qRpL32_f 50 -CGCACCAAGCACTTCATCCG-30; qRpL32_r 50 -GCACGTTGTGCACCAGGAAC-30

**SI Figure Legends**

**Figure S1. Overexpressing Mask in dopaminergic neurons extends lifespan in flies.**

A) Survivorship curves of control flies (Ddc-Gal4 > UAS-tdTomato), flies expressing *mask* RNAi (Ddc-Gal4 > Mask RNAi) or wild type Mask transgenes in the DA neurons (Ddc-Gal4 >UAS- Mask). Knocking down Mask in the DA neurons does not impact the lifespan in the flies, while overexpressing Mask profoundly extend the lifespan in flies.  B) Survivorship curves of control flies (Elav-Gal4 > UAS-tdTomato), flies expressing *mask* RNAi in all neurons (Elav-Gal4 > Mask RNAi), or flies overexpressing Mask in all neurons (Elav-Gal4 > UAS-Mask). Loss-of-function of Mask in all neurons leads to partial lethality during the development (data not shown). Those that do eclose show significantly reduced lifespan. In contrast to the effect of *mask* loss-of-function, overexpressing Mask in all neurons does not affect lifespan in flies.

**Figure S2. The lifespan extension induced by Mask overexpression is specifically mediated by dopaminergic neurons.**

Survivorship curves of flies that overexpress Mask in A) all cells (Da-Gal4, ubiquitous driver); B) glial cells (Repo-Gal4); C) all neurons (elav-Gal4); or D) dopaminergic neurons (Ddc-Gal4). Only when Mask is overexpressed in dopaminergic neurons, the lifespan of the flies is profoundly prolonged. Overexpressing Mask ubiquitously in the entire body or in all neurons does not affect the longevity. Overexpressing Mask in glial cells only moderately extends the lifespan in flies.

**Figure S3. Suppressing the Neuronal Activation of the TH-C’ or TH-D’ Neurons Does not impact the Lifespan in Flies.**

The Survivorship curves of flies expressing the engineered DREADD-Di ($G_{\alpha i}$) in the (A, B) TH-C' or (C, D) TH-D' DANs are shown. The flies are raised on food containing DMSO only (No Drug), 1uM, 10uM or 100uM Clozapine N-oxide (CNO) for their entire adulthood. Inhibiting the activation of the TH-C' or TH-D' DANs does not affect the lifespan in flies.

**Figure S4. Chronically activating the TH-C' and TH-D' dopaminergic neurons in the adult flies moderately shortens lifespan.**

Survivorship curves of male A) and female B) flies expressing the thermal sensitive cation channel TrpA1 in the TH-C' or the TH-D' DANs. The activation of these neurons were achieved by raising the flies at 29°C shortly after eclosion. Long-term activation of the TH-C' neurons leads to 33% reduction in the lifespan (median lifespan shortened from 44 days to 33 days.), and activating the TH-D' neurons results in 10% reduction in the lifespan (median lifespan shortened from 31 days to 28 days.). No effect on lifespan was detected in females when TH-C' or TH-D' dopaminergic neurons were activated by TrpA1.

**Figure S5. Ubiquitously Elevating dopamine activity is not sufficient to extend lifespan in flies.** Survivorship curves of A) female and B) male $w^{1118}$ flies raised on regular food, or on food containing 1mg/ml L-DOPA; and C) female and D) male $w^{1118}$ flies raised on regular food, or on food containing 75uM of SKF-82958, 30uM of Quinpirole, or both drugs are shown. None of the treatments extended the lifespan of male flies. Feeding female flies with dopamine precursor L-DOPA moderately shortened the lifespan (median lifespan reduced from 88 days in the control flies to 78 days in L-DOPA fed flies, ~12% reduction). Neither dopamine receptor agonists alone nor together shows significant effects on female lifespan, although a small population of fed flies showed weaker tolerance and died earlier than untreated group. Log-Rank Test $p$ values: C) $p$ = 0.25 ($w^{1118}$+SKF-82958 vs. $w^{1118}$), $p$ = 0.091 ($w^{1118}$+Quinpirole vs. $w^{1118}$), $p$ = 0.20 ($w^{1118}$+SKF-82958+Quinpirole vs. $w^{1118}$); and D) $p$ = 0.68 ($w^{1118}$+SKF-82958 vs. $w^{1118}$), $p$ = 0.99 ($w^{1118}$+Quinpirole vs. $w^{1118}$), $p$ = 0.29 ($w^{1118}$+SKF-82958+Quinpirole vs. $w^{1118}$).

**Figure S6. Overexpressing Mask in the TH-C' or TH-D' Neurons Does not Change Food Intake in Flies.**

The food intake in a 24 hour interval was measured using a Capillary Feeder assay (CAFE). The amount of food consumed was represented by the distance (mm) between the positions of the liquid food surface at the beginning and the end of the experiments. Food intake of flies that overexpress Mask in the TH-C' neurons is compared to the control flies at A) 10-day or B) 30-day old ages. Food intake of flies that overexpress Mask in the TH-D' neurons is compared to the control flies at C) 10-day or D) 30-day old ages. No differences in food intake was detected between the long-lived flies and the control flies. n=3 for all sample groups.

**Figure S7. Overexpressing Mask in the TH-C' or TH-D' Neurons moderately increase TAG and Glucose levels at mid and late ages in Flies.**

A, B) Age related whole body Glucose/Protein and C, D) TAG/Protein levels in Mask overexpressing or control flies. Mask overexpressing flies contain higher glucose and TAG levels at mid (30-day) and late (50-day) ages, with the exception that female flies overexpressing Mask in the TH-C' neurons show similar levels of TAG compared to the controls. n=4 for all sample groups.

**Figure S8. Overexpressing Mask in the TH-C' or TH-D' DANs does not lead to consistent changes in Dilp2 transcript levels in the adult brains.**

The transcript levels of Dilp2 in the adult brain was measured in newly eclosed and 2-, 5-, 7-, 10-, 15-, 30- and 50-day old flies. A) Male flies overexpressing Mask in the TH-C' neurons show reduced Dilp2 levels at 7- and 15-day old ages. B) Male flies overexpressing Mask in the TH-D' neurons show reduced Dilp2 levels at 50-day old age. C) Female flies overexpressing Mask in the TH-C' neurons show reduced Dilp2 levels at 10-day old age. D) Female flies overexpressing Mask in the TH-D' neurons show no changes in Dilp2 levels in adulthood. *** $P < 0.05$

**Figure S9. Overexpressing Mask in the TH-C' or TH-D' DA neurons does not lead to consistent changes in Dilp3 transcript levels in the adult brains.**

The transcript levels of Dilp3 in the adult brain was measured in newly eclosed and 2-, 5-, 7-, 10-, 15-, 30- and 50-day old flies. A) Male flies overexpressing Mask in the TH-C' neurons show no changes in Dilp3 levels. B) Male flies overexpressing Mask in the TH-D' neurons show reduced Dilp3 levels at 50-day old age. C) Newly eclosed female flies overexpressing Mask in the TH-C' neurons show reduced Dilp3 levels. D) Female flies overexpressing Mask in the TH-D' neurons show reduced Dilp3 levels at 50-day old age. *** $P < 0.05$

**Figure S10. Overexpressing Mask in the TH-C' or TH-D' DA neurons does not lead to consistent changes in Dilp5 transcript levels in the adult brains.**

The transcript levels of Dilp5 in the adult brain was measured in newly eclosed and 2-, 5-, 7-, 10-, 15-, 30- and 50-day old flies. A) Male flies overexpressing Mask in the TH-C' neurons show reduced Dilp5 level at 15-day old age. B) Male flies overexpressing Mask in the TH-D' neurons show reduced Dilp5 levels at 15- and 50-day old ages. C) Female flies overexpressing Mask in the TH-C' neurons show no changes in Dilp5 levels. D) Female flies overexpressing Mask in the TH-D' neurons show no changes in Dilp5 levels. *** $P < 0.05$

**Figure S11. Overexpressing Mask in the TH-C' or TH-D' DA neurons does not lead to consistent changes in Dilp6 transcript levels in the adult brains.**

The transcript levels of Dilp6 in the adult brain was measured in newly eclosed and 2-, 5-, 7-, 10-, 15-, 30- and 50-day old flies. A) Male flies overexpressing Mask in the TH-C' neurons show no changes in Dilp6 levels. B) Male flies overexpressing Mask in the TH-D' neurons show increased Dilp6 level at 30-day old age. At 50-day old age, Dilp6 transcript level in the head is moderately reduced. C) Newly eclosed female flies overexpressing Mask in the TH-C' neurons show reduced Dilp6 level. D) Female flies overexpressing Mask in the TH-D' neurons show reduced Dilp6 level at 50-day old age. *** $P < 0.05$

**Figure S12. The KH domain in the Mask protein is required for Mask to promote autophagy.**

(A) Confocal images of GFP-mCherry auto-fluorescence (using the same gain) from fixed larval muscles in control animals (24B-Gal4>UAS-GFP-mCherry-ATG8), animals overexpressing wild type Mask protein (24B-Gal4>UAS-GFP-mCherry-ATG8+UAS-Mask) or Mask-KH-Mut protein (24B-Gal4>UAS-GFP-mCherry-ATG8+UAS-Mask-KH-Mut). (B) Quantification of mCherry intensity in the larval body wall muscles 6 is shown. Overexpressing Mask in the muscle promotes autophagy, while the KH-Mutant protein fail to induce the same effects.

**Figure S13. Expressing Mask in neurons in the adult flies fail to induce lifespan extension compared to ATG1.**

A, D) Wild type flies and flies B, E) expressing ATG1 (GS-Elav>UAS-ATG1) or C, F) Mask (GS-Elav>UAS-Mask) in their neurons are raised on regular food or food containing 50ug/ml RU488 in order to assess the effects of adulthood neuronal expression of Mask or ATG1 on longevity. GS-Elav induces pan-neuronal expression of transgenes in the presence of RU488. 50ug/ml RU488 induces a moderate adverse effect on lifespan in wild type flies. Expressing ATG1 in neurons leads to moderate lifespan extension (in reference to wild type flies). The lifespan of Mask overexpressing flies, to the contrary, show no difference compared to the non-expressing flies.

**Figure S14. Overexpressing Mask Promotes formation of YFP-Rab5-Positive Membrane Structures at the Presynaptic Terminal at fly larval neuromuscular junctions (NMJs).**

A) Representative confocal images of synapses in control (elav-Gal4>*mask* control RNAi), *mask* knockdown (elav-Gal4>*mask* RNAi) and Mask overexpressing (elav-Gal4>UAS-Mask) larvae are highlighted with HRP (red). YFP auto-fluorescence (using the same gain) from fixed sample marks the YFP-Rab5-positive compartments at the nerve terminals. B) Quantification of the intensity of the YFP signals in the nerve terminal of muscle 6/7 are shown. Knocking down

Mask significantly reduces, while overexpressing Mask, increases the intensity of the YFP-Rab5-positetive compartments.

**Figure S15. Mask Promotes FM1-43 Dye Loading at Fly Larval Neuromuscular Junctions (NMJs).**

A) Representative synapses and boutons in control (elav-Gal4/+) and Mask overexpressing (elav-Gal4 > UAS-Mask) larvae are highlighted with HRP (blue). FM1-43 dye loaded in the vesicle at the nerve terminals is shown in red. B) Quantification of FM1-43 dye intensity in the NMJs on muscle 6/7.

The membrane nonpermeable dye FM1-43 can be taken up into the synaptic vesicles through endocytosis, allowing the quantification of vesicle exocytosis-endocytosis events taking place at the nerve terminals. After stimulation, significantly higher level of FM1-43 loading was detected in presynaptic boutons in Mask overexpressing synapses compared to the controls. No significant FM1-43 loading at the NMJs was detected in a 30 minute interval at rest state in control or Mask overexpressing synapses (data not shown), indicating that overexpressing Mask does not affect endocytosis per se, but rather, the activity-dependent vesicle fusion events.

**Figure S16. Enhancing endocytosis and vesicle trafficking in the TH-C' neurons moderately Extend Lifespan.**

Survivorship curves of flies expressing the wild type Rab5 protein (UAS-YFP-Rab5), the dominant negative form of Rab5 (UAS-YFP-Rab5-S43N), or the constitutively active form of Rab5 protein (UAS-YFP-Rab5-Q88L) are shown. A, B) Overexpressing the wild type Rab5 protein, or C, D) the dominant negative form of Rab5 in the TH-C' neurons does not affect lifespan in flies. E) Overexpressing the constitutively active form of Rab5 protein in the TH-C' neurons moderately extends lifespan in male flies. The median lifespan increased from 68 days in the control flies to 74 days in TH-C'-Gal4 > UAS-YFP-Rab5-Q88L flies, ~8.8% increase. F) Overexpressing the constitutively active form of Rab5 protein in the TH-C' neurons does not affect the lifespan in female flies.

References

1. Zhu, M., Li, X., Tian, X. & Wu, C. Mask loss-of-function rescues mitochondrial impairment and muscle degeneration of Drosophila pink1 and parkin mutants. *Hum Mol Genet* **24**, 3272-3285 (2015). PubMed PMID: 25743185
2. Budnik, V., Koh, Y. H., Guan, B., Hartmann, B., Hough, C., Woods, D. & Gorczyca, M. Regulation of synapse structure and function by the Drosophila tumor suppressor gene dlg. *Neuron* **17**, 627-640 (1996). PubMed PMID: 8893021
3. Kayser, M. S., Yue, Z. & Sehgal, A. A critical period of sleep for development of courtship circuitry and behavior in Drosophila. *Science* **344**, 269-274 (2014). PubMed PMID: 24744368
4. Zhang, M. Z., Yao, B., Wang, S., Fan, X., Wu, G., Yang, H., Yin, H., Yang, S. & Harris, R. C. Intrarenal dopamine deficiency leads to hypertension and decreased longevity in mice. *J Clin Invest* **121**, 2845-2854 (2011). PubMed PMID: 21701066
5. Wiemerslage, L., Schultz, B. J., Ganguly, A. & Lee, D. Selective degeneration of dopaminergic neurons by MPP(+) and its rescue by D2 autoreceptors in Drosophila primary culture. *J Neurochem* **126**, 529-540 (2013). PubMed PMID: 23452092
6. Walters, M. R., Dutertre, M. & Smith, C. L. SKF-82958 is a subtype-selective estrogen receptor-alpha (ERalpha ) agonist that induces functional interactions between ERalpha and AP-1. *J Biol Chem* **277**, 1669-1679 (2002). PubMed PMID: 11700319
7. Diegelmann, S., Jansen, A., Jois, S., Kastenholz, K., Velo Escarcena, L., Strudthoff, N. & Scholz, H. The CApillary FEeder Assay Measures Food Intake in Drosophila melanogaster. *J Vis Exp* (2017). PubMed PMID: 28362419
8. Matsuda, H., Yamada, T., Yoshida, M. & Nishimura, T. Flies without trehalose. *J Biol Chem* **290**, 1244-1255 (2015). PubMed PMID: 25451929

### Supplemental Figure 1

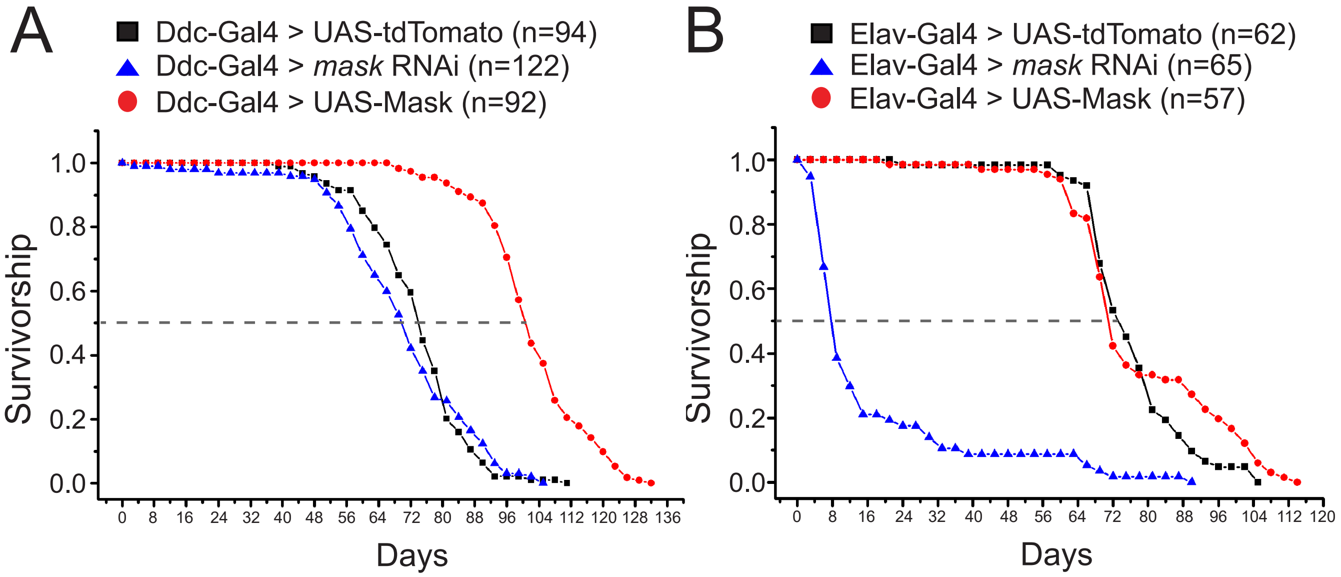


*p < 0.001 Control vs UAS-Mask*

*p = 0.40 Control vs RNAi*

*p < 0.001 Control vs RNAi*

*p = 0.16 Control vs UAS-Mask*

### Supplemental Figure 2

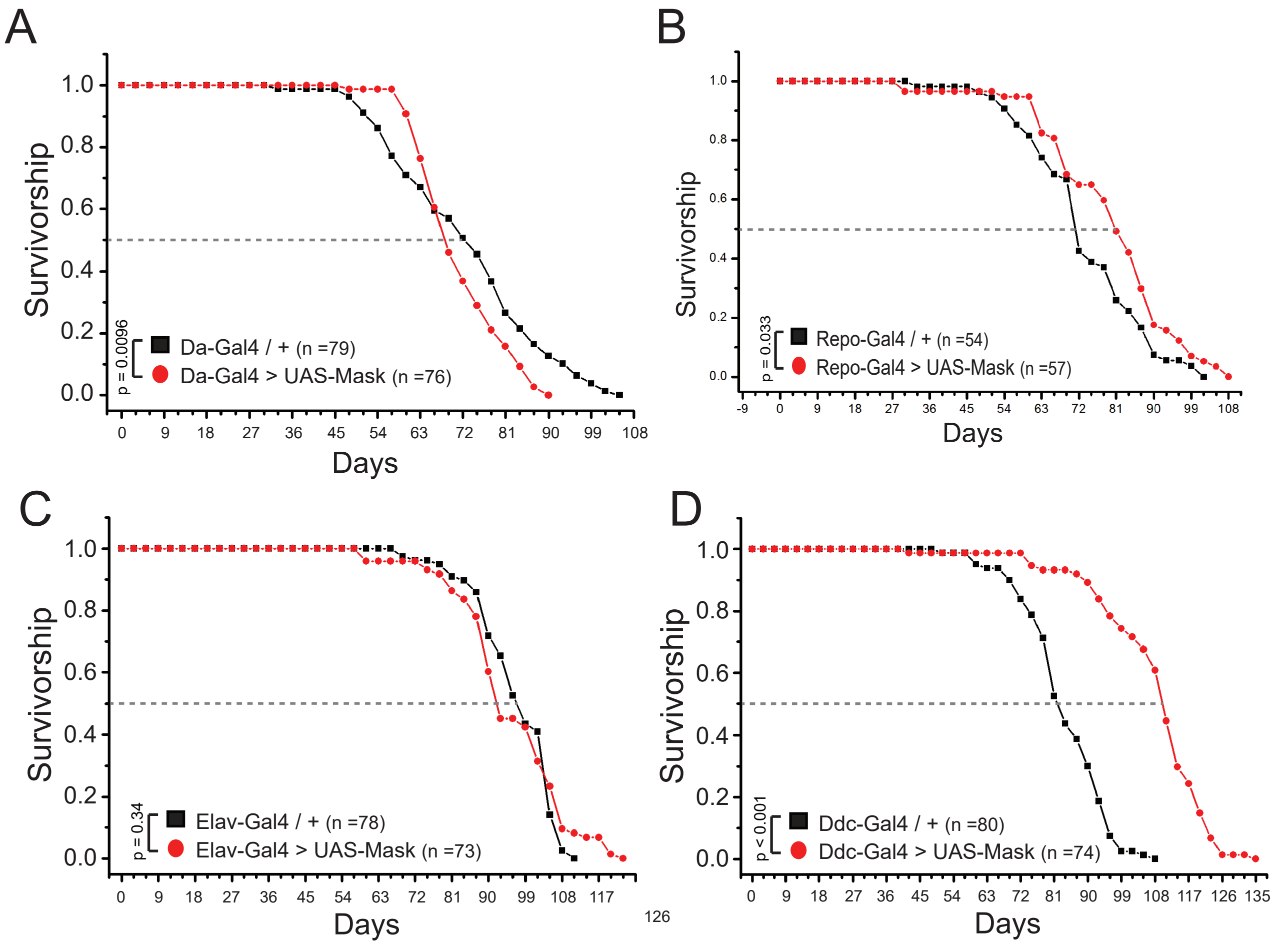

### Supplemental Figure 3

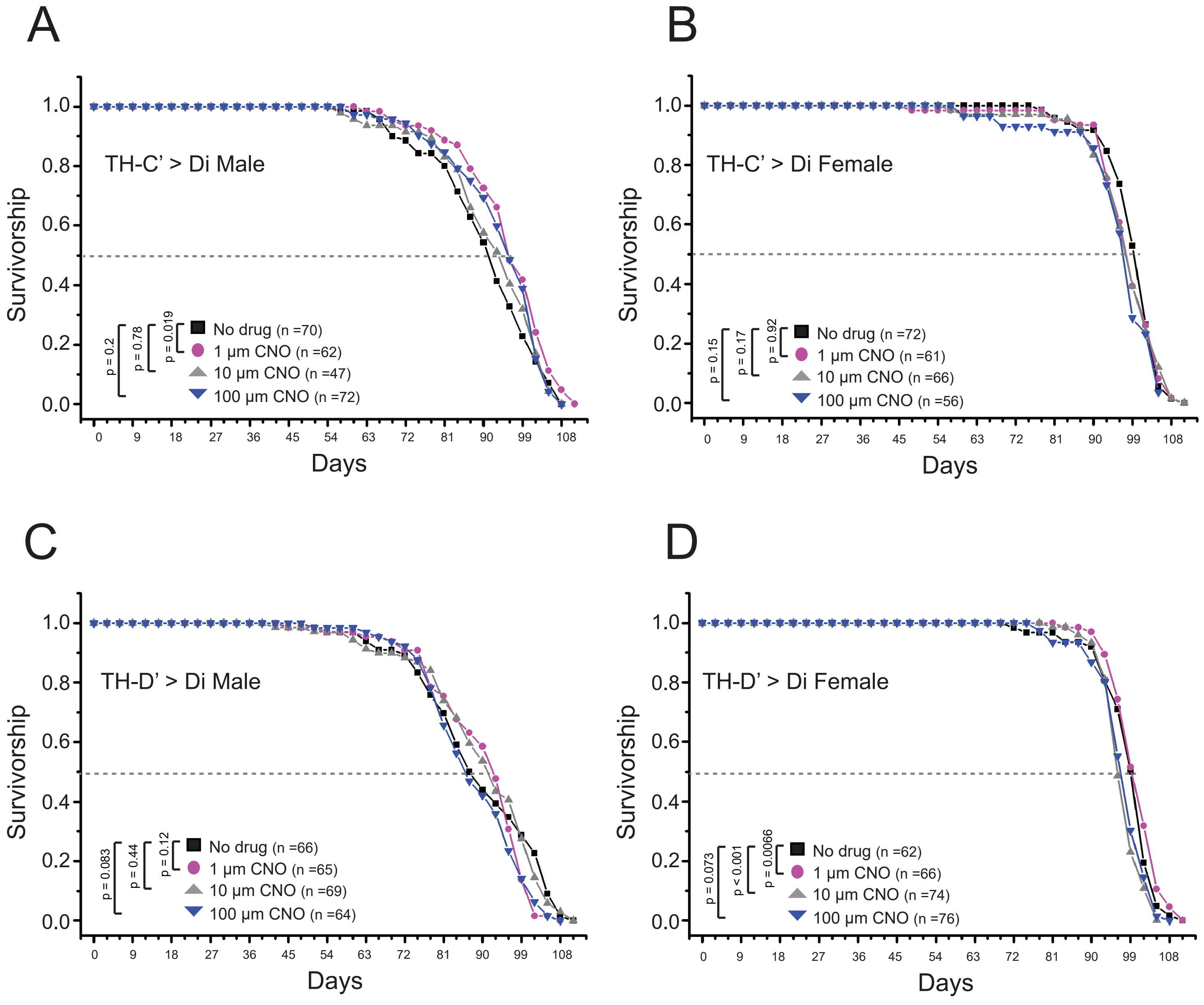

### Supplemental Figure 4

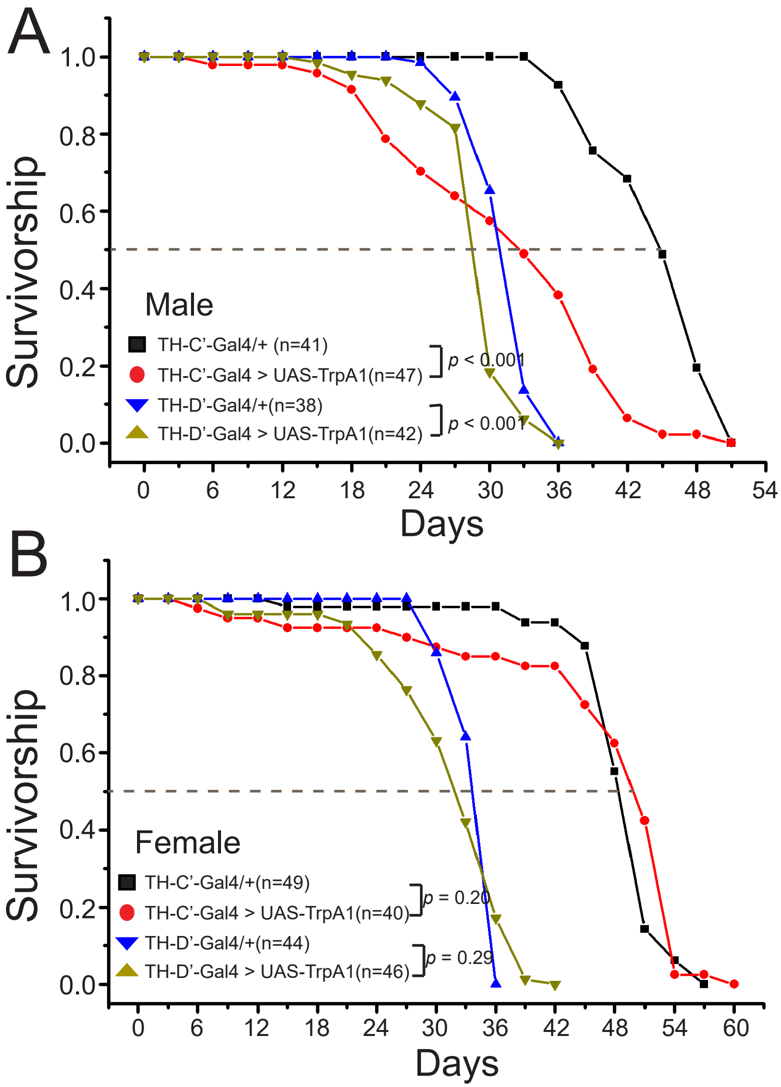

### Supplemental Figure 5

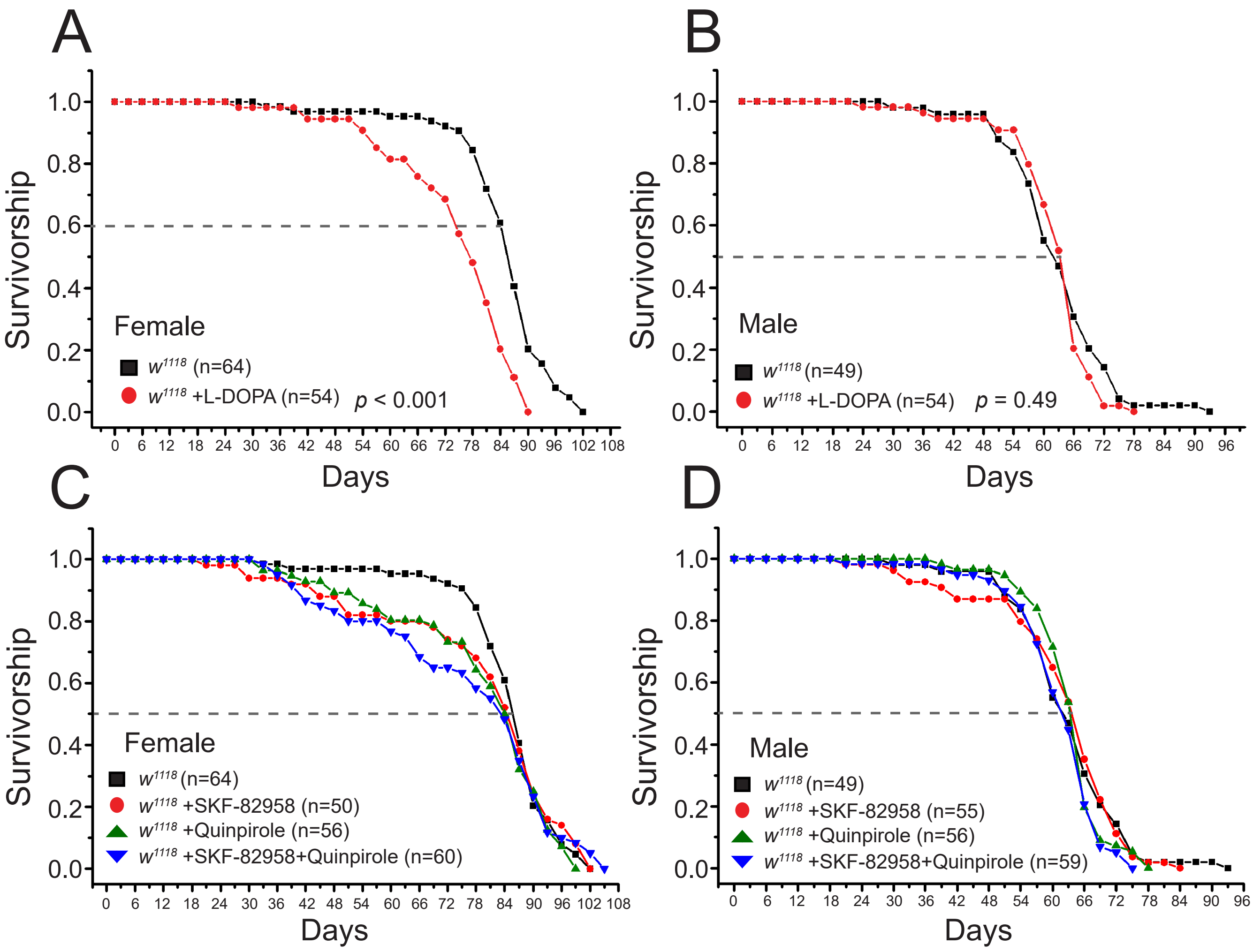

### Supplemental Figure 6

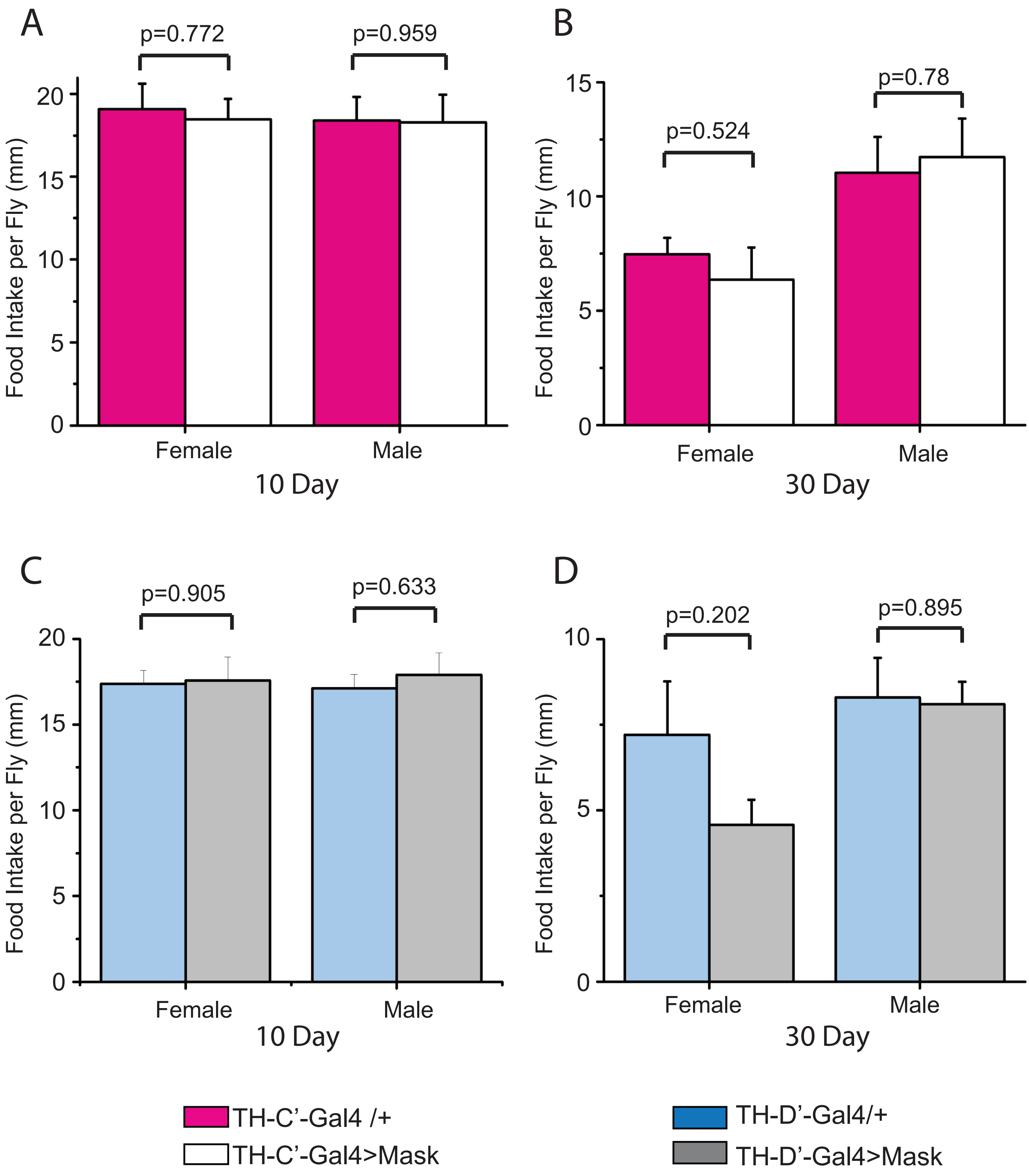

### Supplemental Figure 7

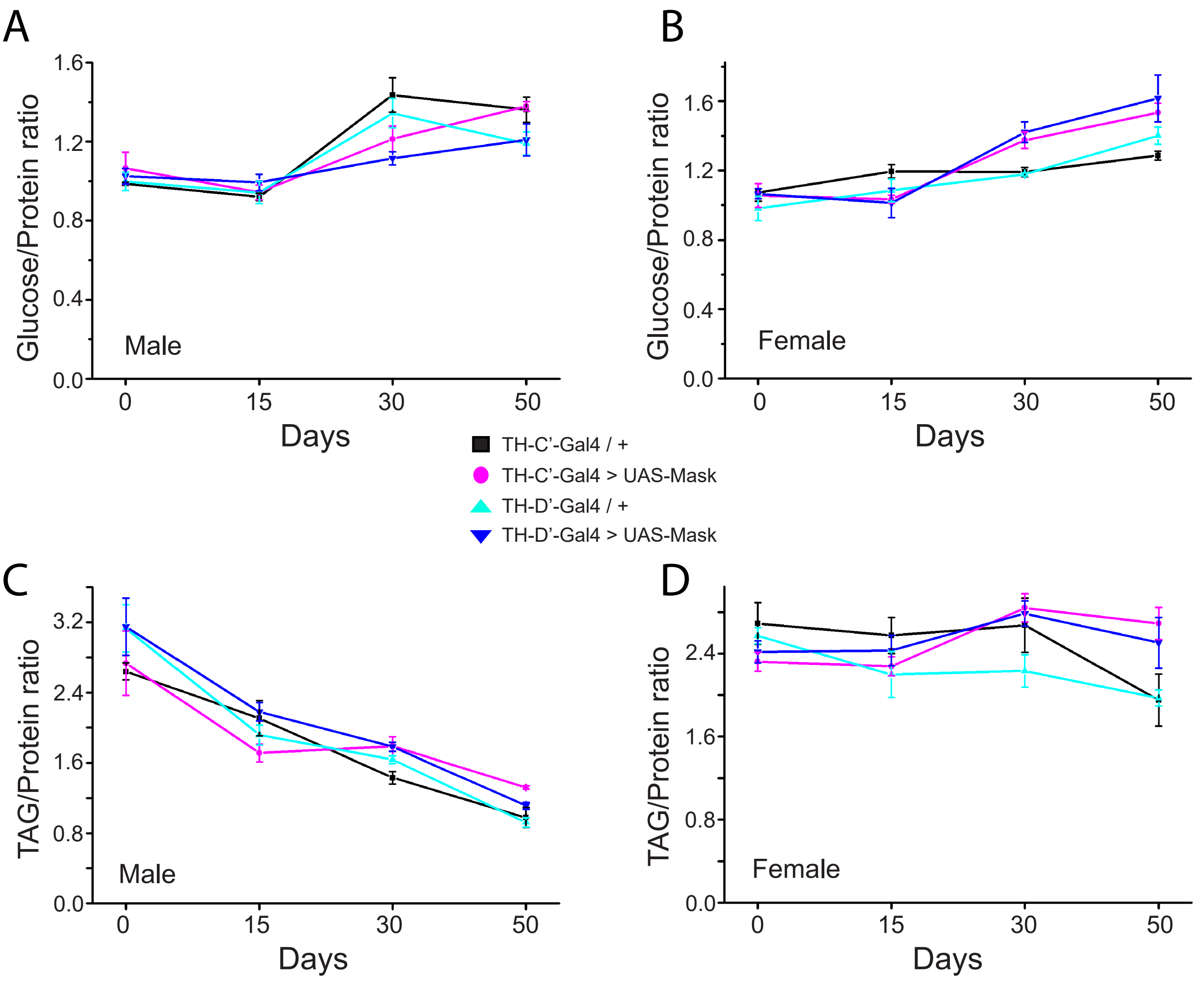

### Supplemental Figure 8

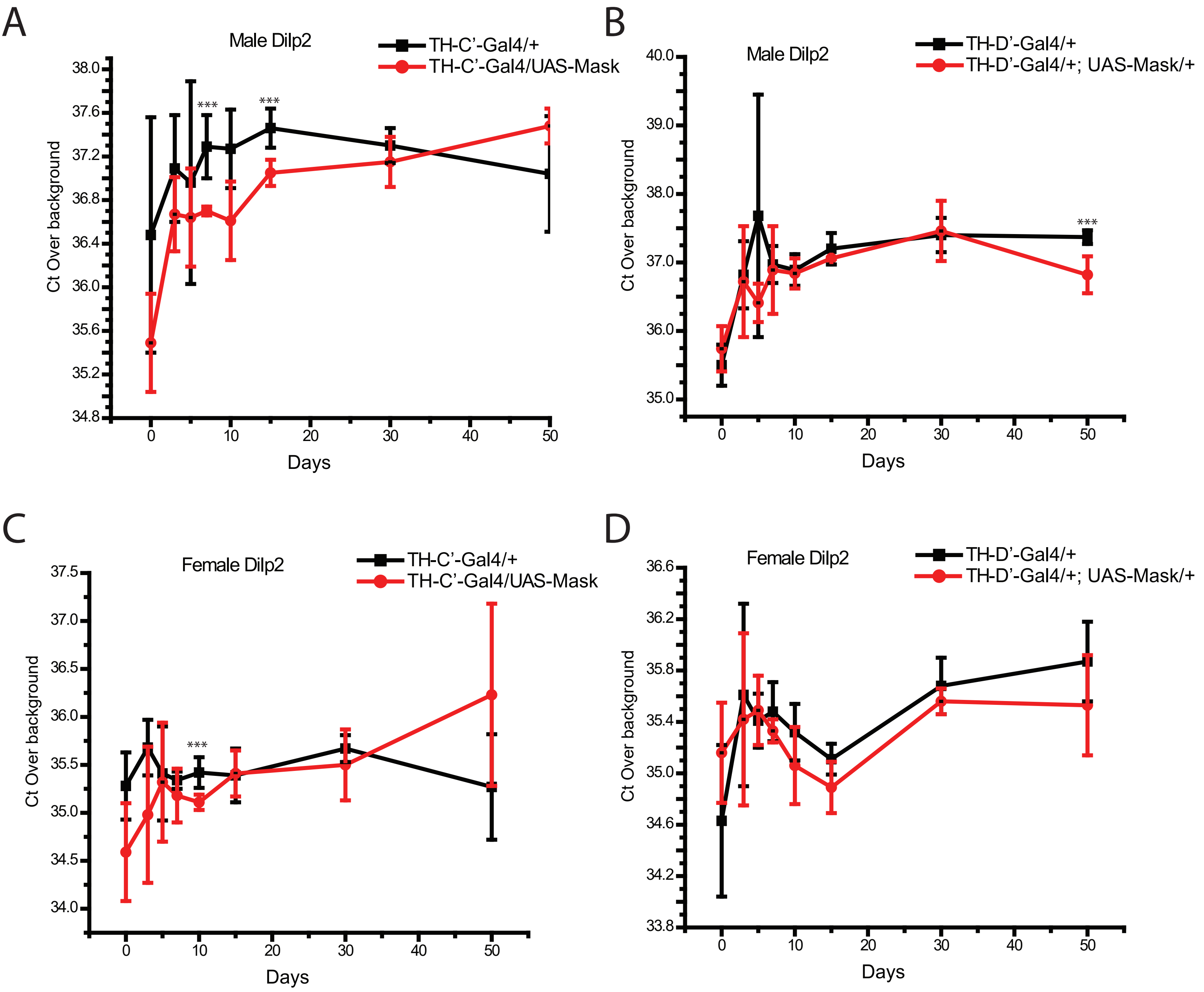

### Supplemental Figure 9

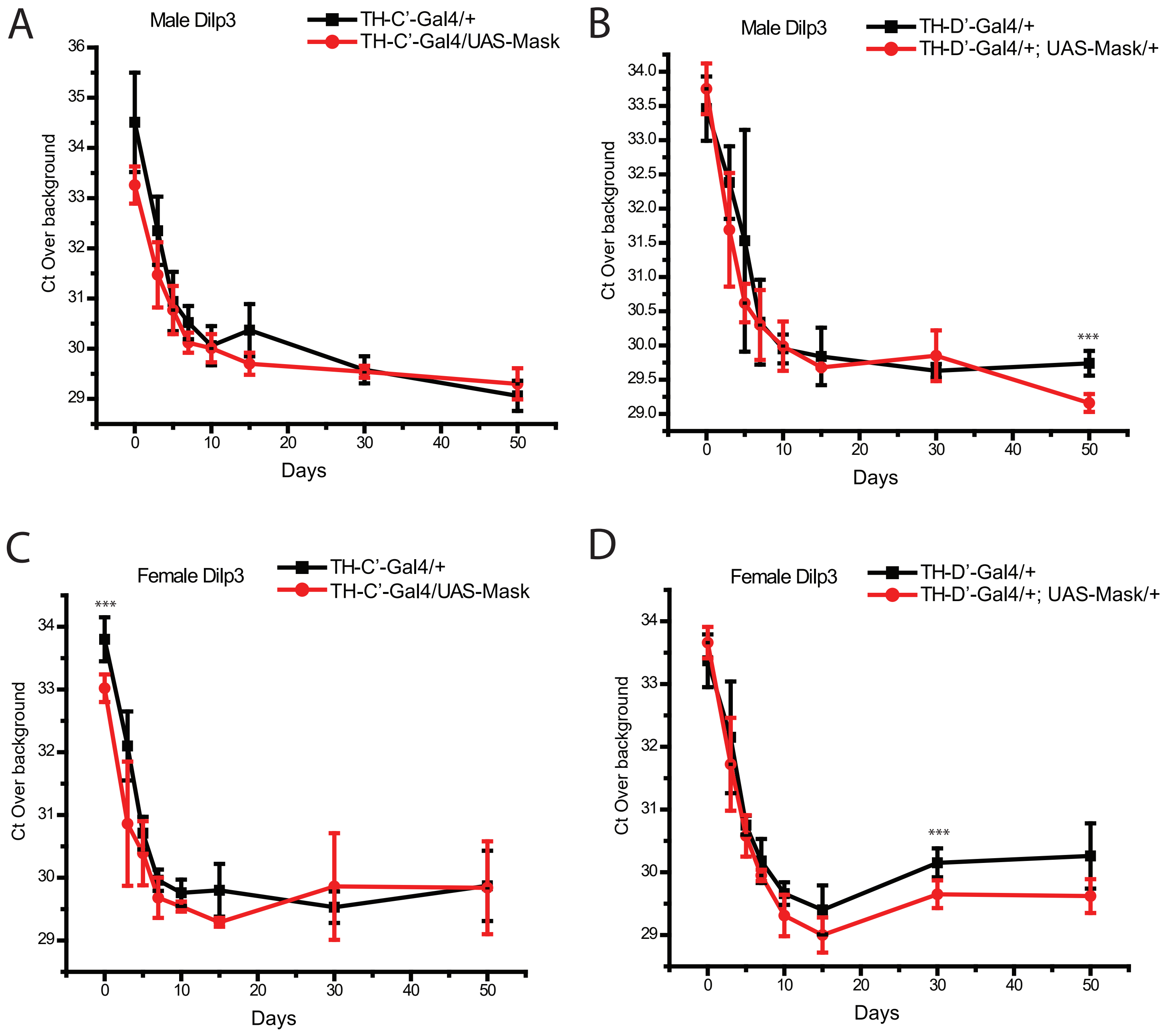

### Supplemental Figure 10

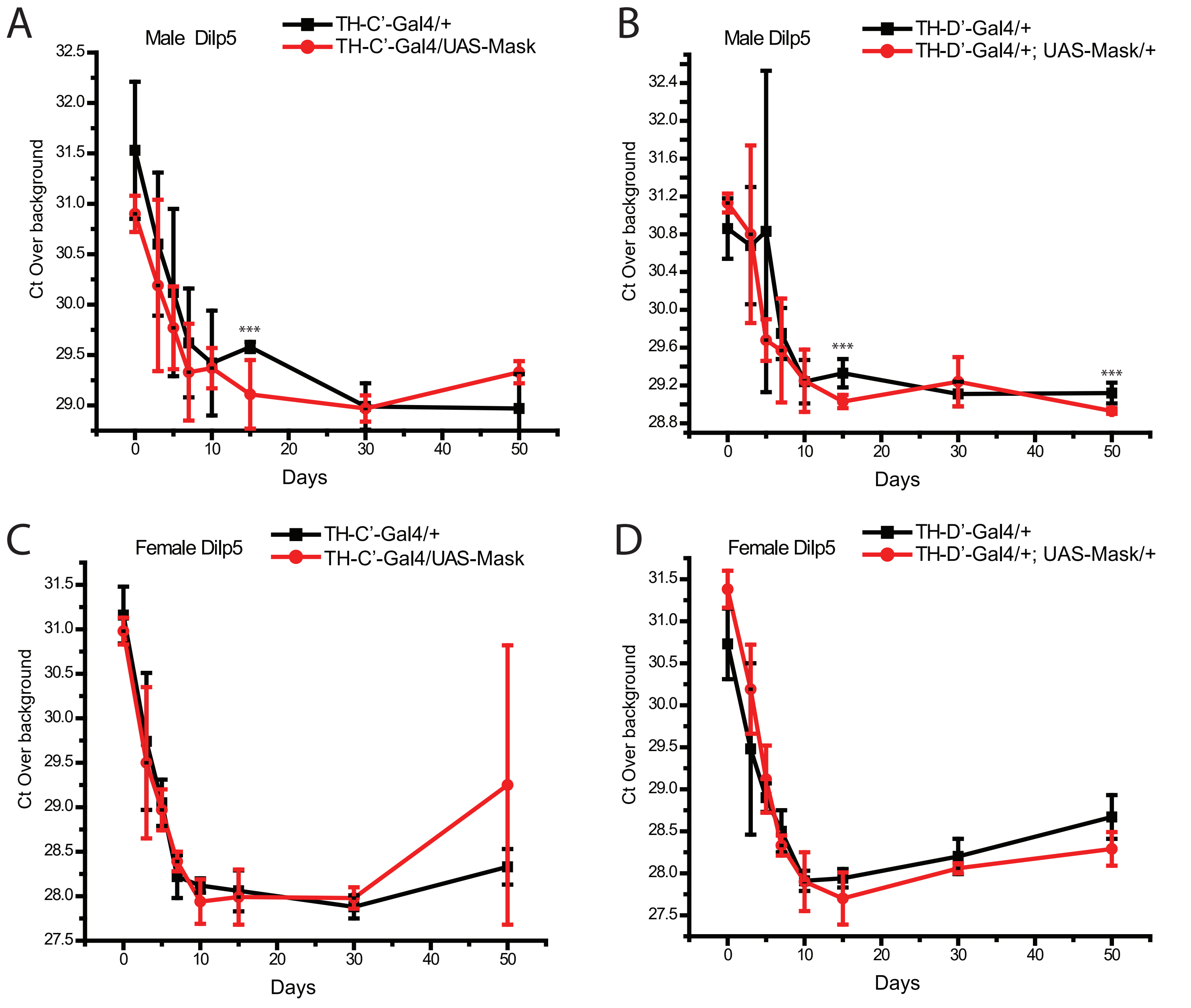

### Supplemental Figure 11

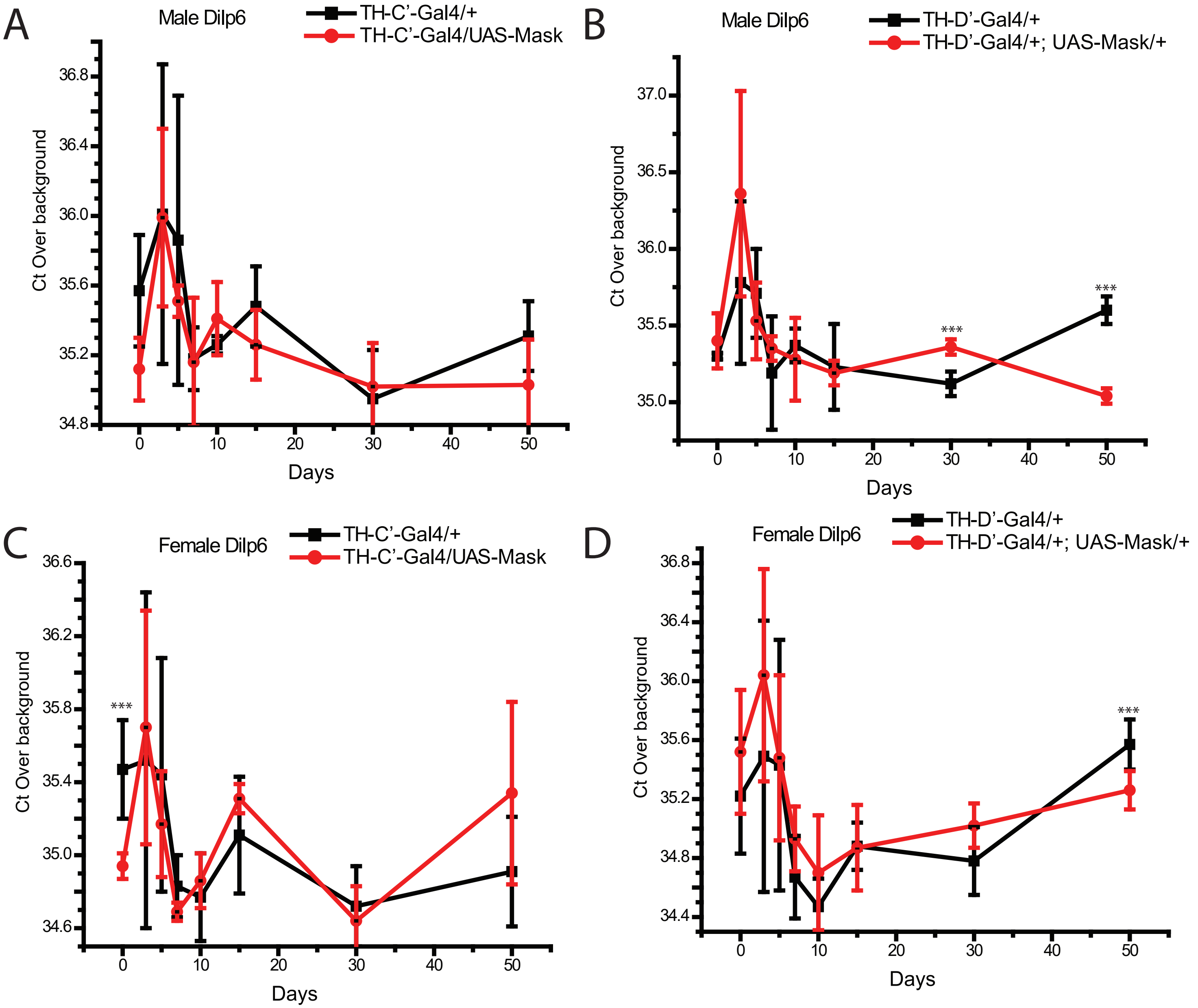

### Supplemental Figure 12

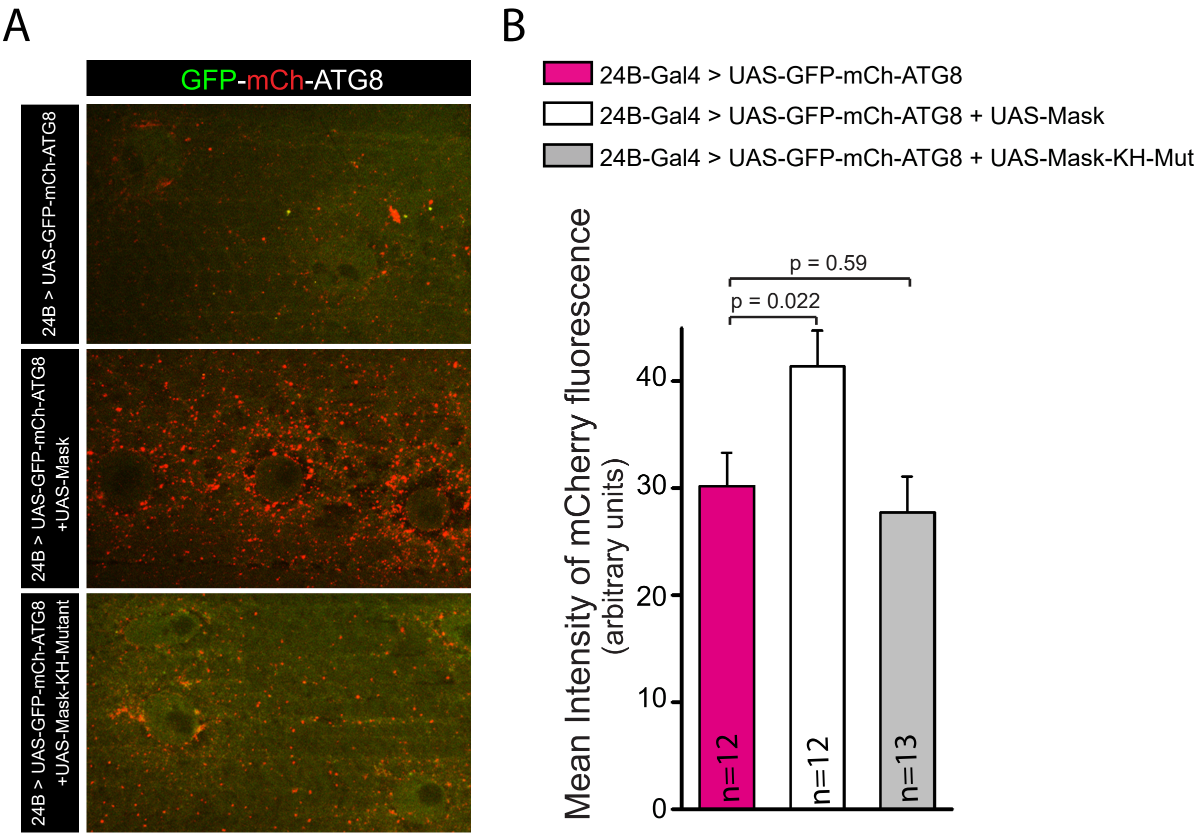

### Supplemental Figure 13

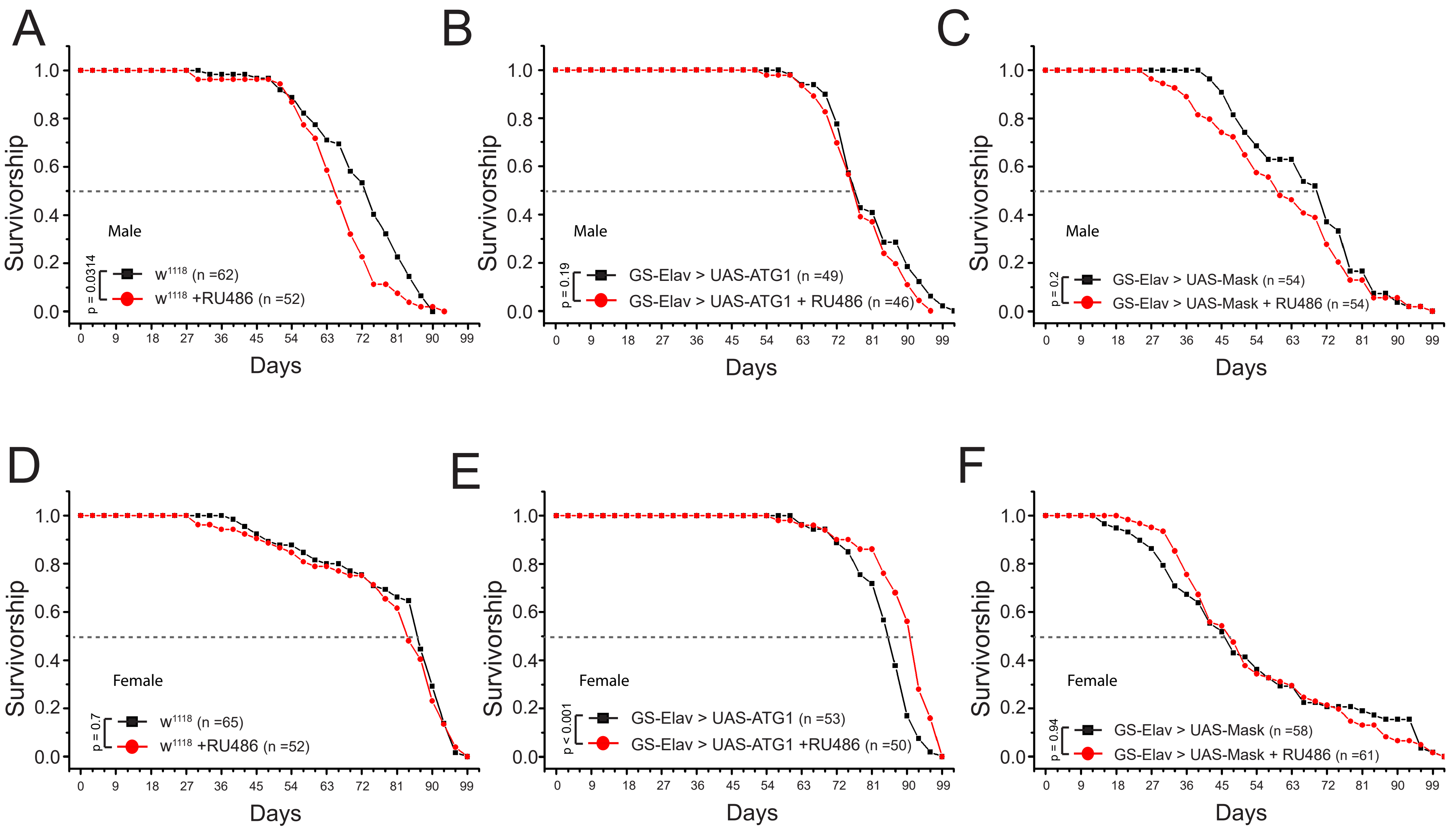

### Supplemental Figure 14

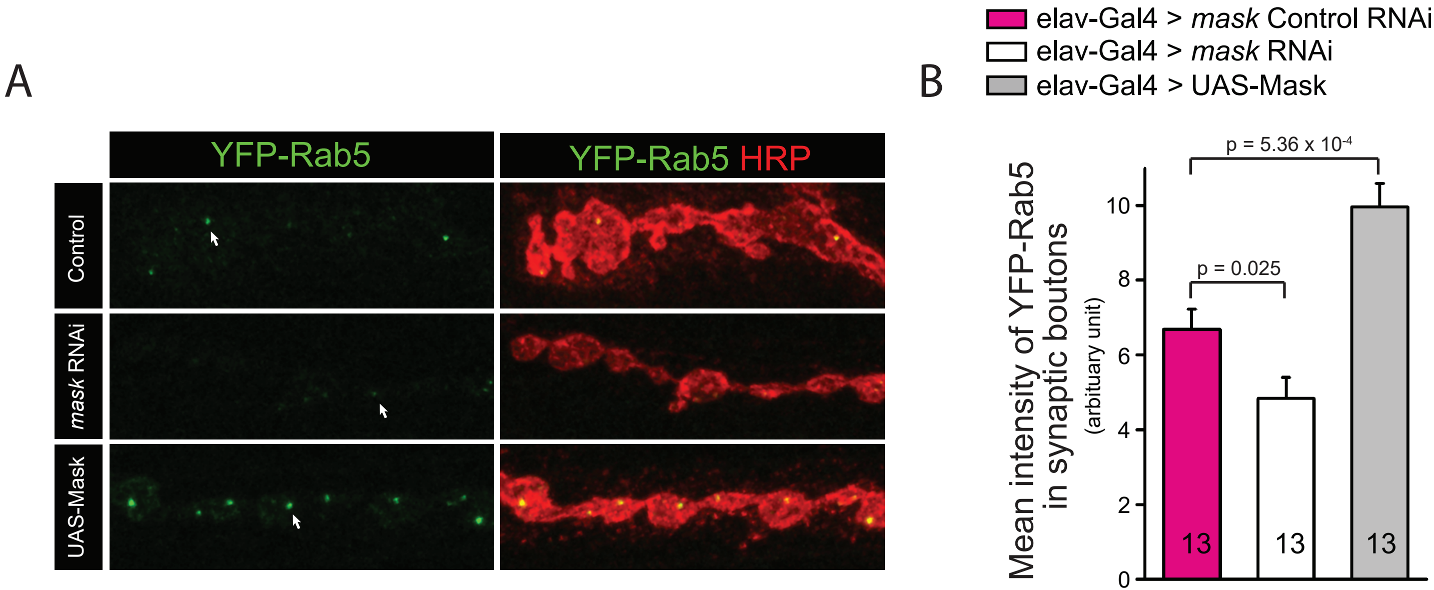

### Supplemental Figure 15

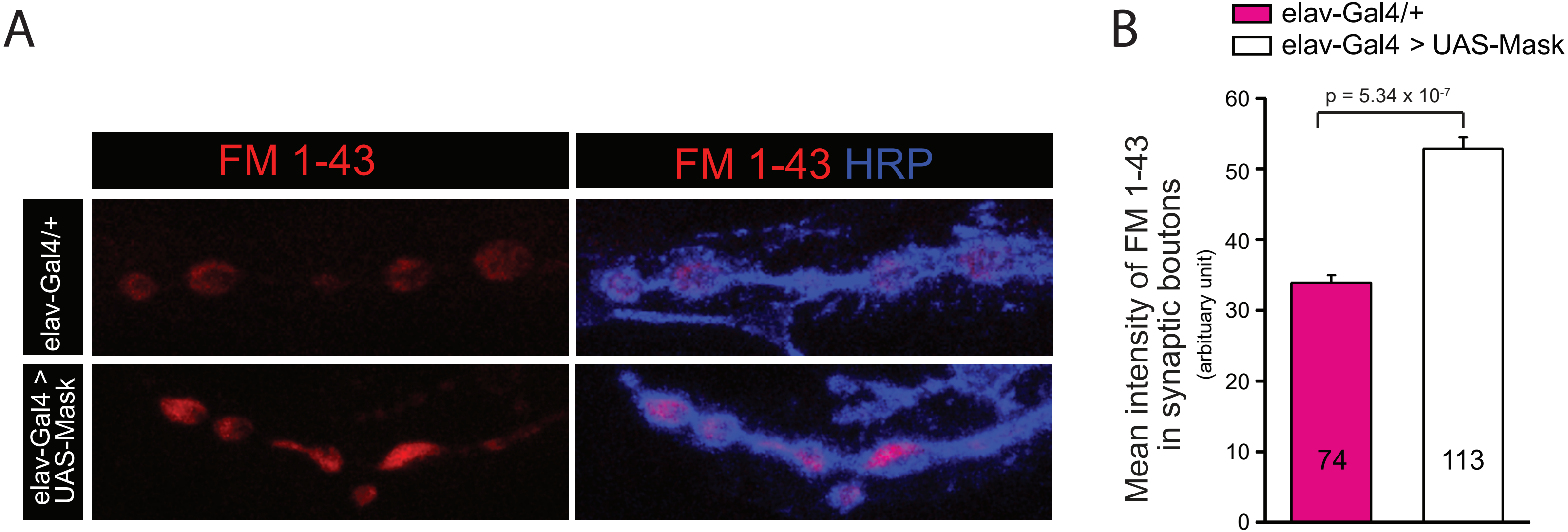

### Supplemental Figure 16

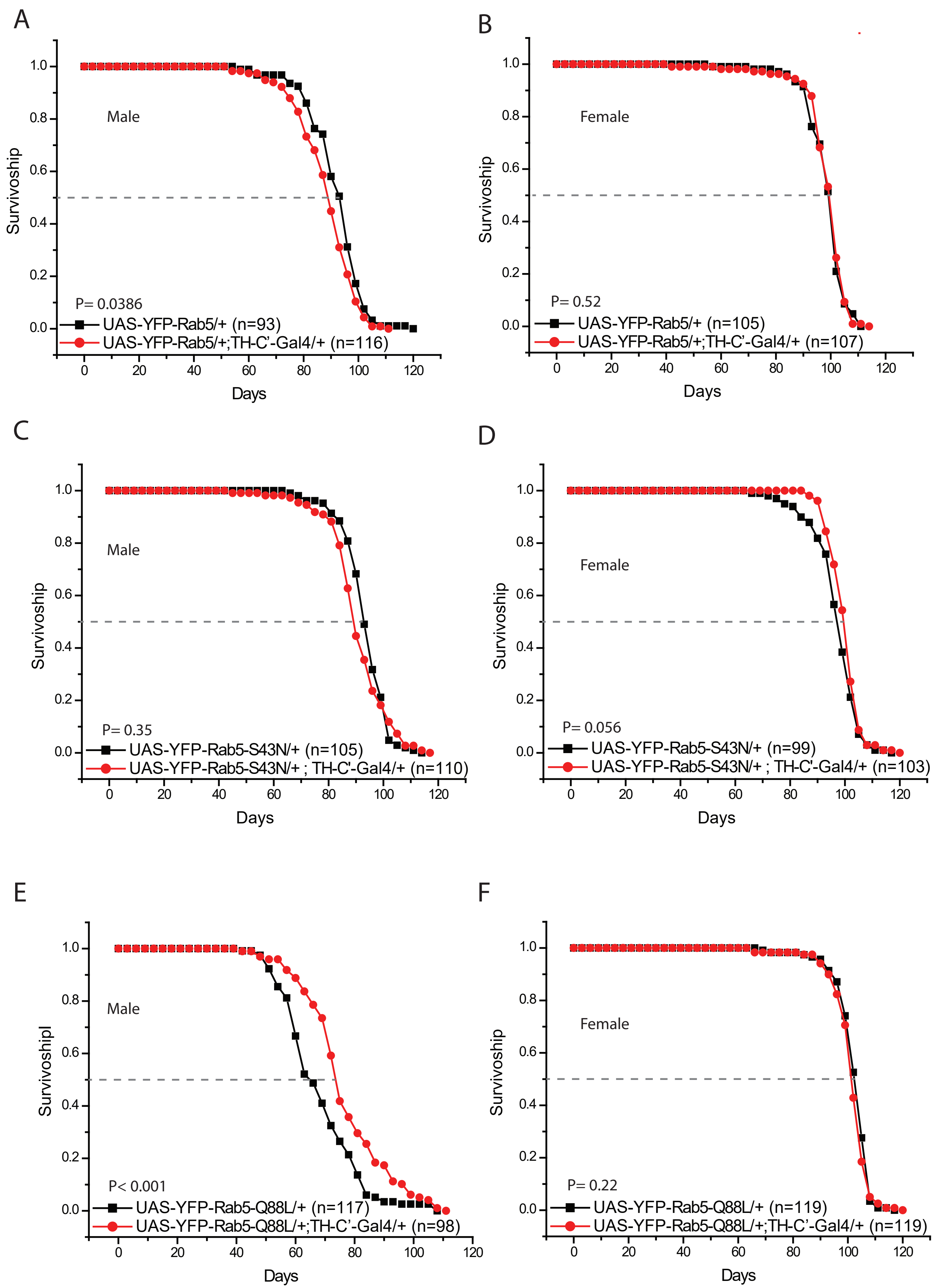
